# A multicolor suite for deciphering population coding in calcium and cAMP *in vivo*

**DOI:** 10.1101/2023.01.06.522686

**Authors:** Tatsushi Yokoyama, Satoshi Manita, Hiroyuki Uwamori, Mio Tajiri, Itaru Imayoshi, Sho Yagishita, Masanori Murayama, Kazuo Kitamura, Masayuki Sakamoto

## Abstract

cAMP is a pivotal second messenger regulated by various upstream pathways including Ca^2+^ and G protein-coupled receptors (GPCRs). To decipher *in vivo* cAMP dynamics, we rationally designed cAMPinG1, an ultrasensitive genetically encoded green cAMP indicator that outperformed its predecessors in both dynamic range and cAMP affinity. Two-photon cAMPinG1 imaging detected cAMP transients in the somata and dendritic spines of neurons in the mouse visual cortex on the order of tens of seconds. In addition, multicolor imaging with a highly sensitive new red Ca^2+^ indicator RCaMP3 allowed simultaneous measurement of population patterns in Ca^2+^ and cAMP in hundreds of neurons. We identified Ca^2+^-induced cAMP responses that represented specific information, such as direction selectivity in vision and locomotion, as well as GPCR-induced cAMP responses. Overall, our multicolor suite revealed that information encoded in Ca^2+^ and GPCRs signaling is integrated and stored as cAMP transients for longer periods *in vivo*.

**Highlights:**

- Developing an ultrasensitive cAMP indicator, cAMPinG1, for visualizing cAMP transients in somata and dendritic spines *in vivo*.
- Developing a highly sensitive red Ca^2+^ indicator, RCaMP3, for visualizing Ca^2+^ transients in large neuronal population.
- Dual-color Ca^2+^ and cAMP imaging for dissecting Ca^2+^-induced and GPCR-induced cAMP responses.
- Single-cell, single-timepoint cAMP imaging for GRCR biology and drug screening.

## INTRODUCTION

Cyclic adenosine monophosphate (cAMP) is a pivotal second messenger that plays a universal role in intracellular signal transduction in a variety of cell types and organisms. Individual cell types express various adenylate cyclases (ACs) and phosphodiesterases (PDEs) that synthesize and degrade cAMP, respectively. The upstream regulators of the ACs are generally G protein-coupled receptors (GPCRs), which increase or decrease intracellular cAMP in a cell-type-specific manner. Some cell types, including neurons, also express ACs that are dependent on another central second messenger, Ca^2+^ (Kandel et al., 2014). The regulation of cAMP by these multiple upstream signaling pathways occurs continuously in the soma and small cellular compartments, such as dendritic spines and axonal boutons, modulating diverse cellular functions through cAMP-dependent kinases, channels, and transcription factors. Despite extensive knowledge of cAMP functions as a second messenger, the precise timing and location of its regulatory effects *in vivo* are still unknown. Therefore, technologies to visualize the spatiotemporal dynamics of cAMP *in vivo* are crucial for biological research in various organs or species.

Since cAMP was first visualized in 1991, more than 50 cAMP indicators have been developed (Adams et al., 1991; Massengill et al., 2021). The circularly permuted green fluorescent protein (cpGFP)-type cAMP indicators have been intensively developed more recently due to their large dynamic range (Kawata et al., 2022; Liu et al., 2022; Wang et al., 2022). However, the use of these indicators has been limited because they have low cAMP affinity (> 1 μM). Since the cAMP affinity of endogenous cAMP-dependent kinases and channels is typically in the hundreds of nanomolar range (Ludwig et al., 1998; Zhang et al., 2012), it is critical to have a submicromolar affinity to detect bidirectional cAMP change *in vivo*. Due to the lack of cAMP indicators with both large dynamic range and submicromolar cAMP affinity, the basic properties of *in vivo* cAMP dynamics remain unclear. Specifically, the following questions have not been answered: (1) What is the time scale of cAMP dynamics in individual cells and subcellular compartments? (2) What information is encoded in the cAMP population pattern? (3) How do multiple upstream signals, such as neuromodulators, GPCRs, and Ca^2+^, influence the population pattern of cAMP change?

Here, we present cAMPinG1, a green cAMP indicator with a significantly larger dynamic range and more than 4.8-fold higher cAMP affinity than the existing green cAMP indicators. *In vivo* cAMP imaging with cellular and subcellular resolution revealed that cAMP transients occurred on the temporal scale of seconds to tens of seconds. We also introduce RCaMP3, an improved red calcium indicator. Dual-color imaging for Ca^2+^ and cAMP revealed that cell-specific cAMP transients represented specific information as a downstream of Ca^2+^ signaling as well as GPCR-induced cAMP responses. In addition, the combination of cAMPinG1 and red-shifted channelrhodopsin (ChRmine) revealed that action potentials are sufficient to induce cAMP transients *in vivo*. Overall, our multicolor suite for Ca^2+^ and cAMP imaging allows us to examine how the information encoded in action potentials, Ca^2+^, and GPCR signaling is integrated and stored for a longer timescale as cAMP transients. We also demonstrated the application of cAMPinG1 imaging in cultured cells for GPCR biology and drug screening.

## RESULTS

### Rational engineering of an ultrasensitive cAMP sensor

To develop a high affinity cAMP sensor, we chose a mammalian protein kinase A regulatory subunit (PKA-R) as the cAMP sensing domain. PKA-Rs are widely distributed and functional in mammalian neurons even when tagged by GFP and overexpressed (Zhong et al., 2009) and have been well characterized in terms of their biochemical, evolutionary, and structural properties (Canaves and Taylor, 2002; Kim et al., 2007; Su et al., 1995; Zhang et al., 2012). We used the cAMP-binding domain A of mammalian PKA-R type 1α (PKA-R1α) because of its high affinity for cAMP (around 150 nM) (Lorenz et al., 2017), which is within the range of the affinity of cAMP-binding domains in multiple PKAs and cyclic nucleotide-gated ion channels (Ludwig et al., 1998; Zhang et al., 2012) (**Figure S1A**). We then inserted cpGFP into the β4 – β5 loop of PKA-R1α for the following reasons: (1) it is close to cAMP in the cAMP-bound three-dimensional structure, (2) it is exposed on the surface, and (3) it is structurally flexible, as demonstrated by the analyses of crystallization and evolutionarily conserved sequences (Canaves and Taylor, 2002; Wu et al., 2004) (**Figure 1A; Figures S1B and S1C**). The structural flexibility of loops was critical to avoid possible structural perturbations that could decrease the affinity (Dagliyan et al., 2016). In addition, we removed the N-terminal PKA-R1α region, which includes a dimerization/docking domain interacting with scaffold proteins and an inhibitory domain interacting with PKA catalytic subunits (PKA-C), to avoid interaction with these endogenous proteins. Instead, we fused the RSET sequence to the N-terminus of the sensor to promote stable expression (Wu et al., 2004). To develop this construct with larger ΔF/F, we then generated a library of over 250 mutants with mutations on the putative interface between the cpGFP and cAMP-binding domain, including two linkers and residues in cpGFP close to the interface, and screened them in *Escherichia coli* (*E. coli*). The variant with the largest fluorescence response to cAMP was named cAMPinG1 (<u>cAMP in</u>dicator <u>G</u>reen <u>1</u>) (**Figure 1B**). Biophysical characterization showed that cAMP-free cAMPinG1 had a dominant excitation peak at 400 nm and a second peak at 516 nm, while cAMP-saturated cAMPinG1 had a dominant excitation peak at 498 nm (**Figure 1C**). The green fluorescence intensity of cAMPinG1 increased by 1,000 % upon binding to cAMP with blue light (488 nm), while the green fluorescence intensity decreased by 61 % with violet light (405 nm), resulting in a 2,700 % ratio change with a combination of blue and violet excitation in HEK293T cell lysate. The large ratio change dependent on the decrease of fluorescence intensity with violet excitation indicates that cAMPinG1 is suitable for ratiometric imaging, in contrast to some cpGFP-type indicators such as GCaMPs and G-Flamp1, which do not show a fluorescence decrease with violet excitation (Inoue et al., 2019; Wang et al., 2022) (**Figure S2A**).

**Figure 1.**
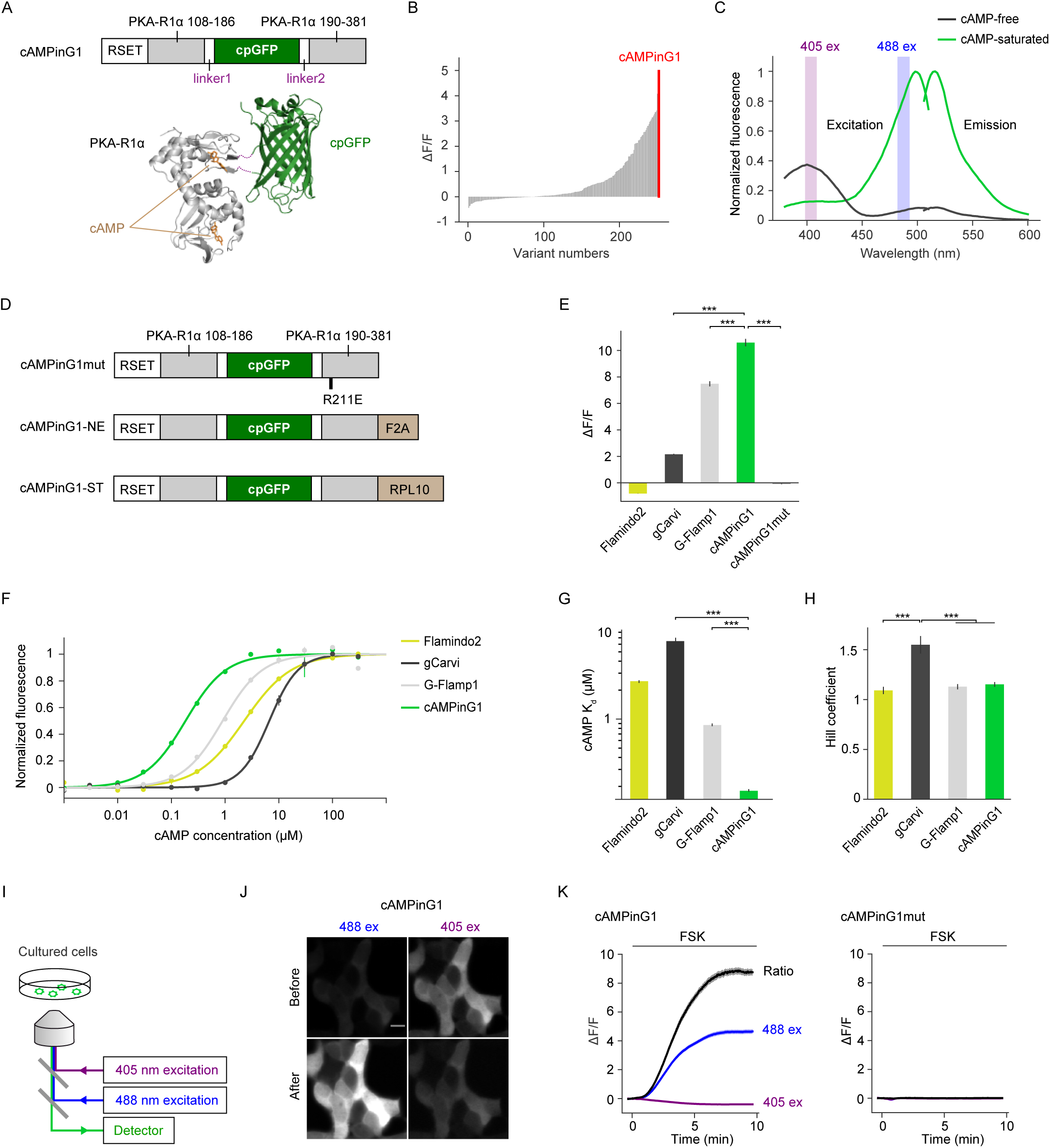
Sensor design and *in vitro* characterization of cAMPinG1. **(A)** Top: Primary structure of cAMPinG1. cpGFP and two flanking linkers are inserted into a loop of PKA-R1α close to the cAMP-binding site. Bottom: Tertiary structures of cAMP-binding PKA-R1α (gray, PDB: 1RGS, cpGFP insertion site of cAMPinG1 is hidden) with cAMP (orange) and cpGFP (green, PDB: 3WLD, calmodulin, and M13 domains are hidden) are shown. The linkers between the two domains are depicted as purple dotted lines. **(B)** *In vitro* screening results of 251 variants, resulting in the variant named cAMPinG1 (red). **(C)** Excitation and emission spectra of cAMPinG1 in cAMP-free (black) and cAMP-saturated (light green) states. Blue and violet excitation wavelengths used for ratiometric imaging in the later figures are indicated as shaded regions in the later figures. Note that green fluorescence intensity increased upon cAMP binding when excited with blue light (488 nm) but decreased when excited with violet light (405nm). **(D)** Primary structures of cAMPinG1mut (inactive mutant), cAMPinG1-NE (nuclear-excluded), and cAMPinG1-NE (soma-targeted). **(E)** The change in fluorescence intensity (ΔF/F) of green cAMP sensors to cAMP in HEK cell lysate. cAMPinG1 had the largest ΔF/F in the side-by-side comparison. n = 4 wells (Flamindo2), n = 4 wells (gCarvi), n = 4 wells (G-Flamp1), n = 4 wells (cAMPinG1), n = 4 wells (cAMPinG1mut). Tukey’s post hoc test following one-way ANOVA. **(F)** cAMP titration curves of cAMP sensors. Response of Flamindo2 to cAMP is inversed (-ΔF/F). The x-axis is logarithmic. n = 4 wells (Flamindo2), n = 4 wells (gCarvi), n = 4 wells (G-Flamp1), n = 4 wells (cAMPinG1). (G) K_d_ values of cAMP sensors. cAMPinG1 and cAMPinG1-NE had the highest cAMP affinity among the green cAMP sensors. The y-axis is logarithmic. n = 4 wells (Flamindo2), n = 4 wells (gCarvi), n = 4 wells (G-Flamp1), n = 4 wells (cAMPinG1). Tukey’s post hoc test following one-way ANOVA. **(H)** Hill coefficients of cAMP sensors. n = 4 wells (Flamindo2), n = 4 wells (gCarvi), n = 4 wells (G-Flamp1), n = 4 wells (cAMPinG1). Tukey’s post hoc test following one-way ANOVA. **(I)** Schematic of the imaging settings. Blue (488 nm) and violet (405 nm) excitation lights were used in turns for ratiometric imaging in HEK293T cells. **(J)** Representative images of HEK293T cells expressing cAMPinG1 excited by blue (488 nm, left) and violet (405 nm, right) lights before (top) and after (bottom) 50 μM forskolin application. Scale bar, 10 μm. **(K)** ΔF/F of cAMPinG1 (left) and the inactive mutant cAMPinG1mut (right) in response to 50 μM forskolin application. Blue (blue line) and violet lights (violet line) were used for excitation sequentially, and the ratio of fluorescence excited with blue and violet lights (488 ex / 405 ex) was also shown (black line). n = 196 (cAMPinG1), n = 164 (cAMPinG1mut) cells. All shaded areas and error bars denote the SEM.

We next developed a cAMP-insensitive indicator (cAMPinG1mut) by introducing R211E (in the numbering of mouse PKA-R1α) mutation to block cAMP binding (**Figure 1D**). cAMPinG1 is distributed throughout the cell, including the nucleus and cytoplasm, considerably affecting signal detection. To detect cAMP changes selectively in the cytoplasm, we added a self-cleaving peptide (F2A), known to work as a nuclear export signal (Ohkura et al., 2012), to the C-termini of cAMPinG1 (named cAMPinG1-NE) (**Figure 1D**). Furthermore, to avoid contamination of somatic neuropil fluorescence signals, we linked ribosomal subunit protein (RPL10) to the C-terminal of cAMPinG1 as soma targeting (named AMPinG1-ST) (Chen et al., 2020) (**Figure 1D**).

### *In vitro* characterization of cAMPinG1

We next investigated side by side several recently developed cpGFP-type cAMP sensors. The comparative evaluation revealed that cAMPinG1 had the largest dynamic range (ΔF/F) and fluorescence intensity in a cAMP-saturated state in HEK293T cell lysate than the existing cAMP sensors, Flamindo2, gCarvi, and G-Flamp1 (Hackley et al., 2018; Kawata et al., 2022; Odaka et al., 2014) (**Figure 1E; Figure S2B; Table S1**). cAMPinG1mut did not respond to cAMP, indicating that the fluorescence change depends on cAMP binding. We then compared the cAMP affinities of these sensors. The concentration-response curve showed that the K_d_ value of cAMPinG1 was 181 nM, less than a quarter of those of Flamindo2, gCarvi, and G-Flamp1(**Figures 1F and 1G**). Furthermore, the cAMP binding to cAMPinG1 was accompanied by Hill coefficients close to 1, indicating the linear relation of cAMP concentration and cAMPinG1 fluorescent intensity (**Figure 1H**). cAMPinG1 showed lower affinity to cyclic guanosine monophosphate (cGMP) (K_d_ = 12 µM), 60-fold larger than the cAMP K_d_ value of cAMPinG1, indicating cAMPinG1 fluorescence change was specific for cAMP (**Figure S2C**).

To further demonstrate the sensitivity in live cell imaging, we transiently expressed cAMPinG1in HEK293T cells by lipofection and performed time-lapse imaging. We applied forskolin, an activator of ACs, to generate a high level of cAMP and assess the maximum fluorescence change. As previously shown (**Figure 1C**), cAMPinG1 is capable of ratiometric imaging by using alternating blue (488 nm) and violet (405 nm) excitation, which can reduce motion artifacts, sensor concentration changes, and ambient light. As expected, the fluorescence intensity increased with 488 nm excitation and decreased with 405 nm excitation in response to the stimulus. While cAMPinG1 had high ΔF/F (∼400%) enough for intensiometric measurement with 488 nm excitation, ratiometric measurement had even higher ΔF/F (∼800%). We also found that cAMPinG1 binding to PKA-C was undetectable (**Figure S2D**). The elevation of cAMPinG1 fluorescence by forskolin administration was also observed in neurons in acute brain slices (**Figure S3**).

### *In vivo* two-photon imaging of cAMP dynamics with subcellular resolution

To demonstrate *in vivo* functionality of cAMPinG1, we introduced cAMPinG1 or inactive cAMPinG1mut into pyramidal neurons in layer 2/3 (L2/3) of the mouse primary visual cortex (V1) by *in utero* electroporation (**Figure 2A**). We applied an aversive airpuff stimulus in the awake condition to induce an elevation of noradrenaline and cAMP in the cortex (Oe et al., 2020). Somatic cAMPinG1 imaging visualized cAMP transients induced by 20 seconds of airpuff with single-cell resolution (**Figures 2B-2E**). Inactive cAMPinG1mut imaging did not show a fluorescence change in response to airpuff, indicating cAMPinG1 fluorescence change was dependent on cAMP binding to cAMPinG1 *in vivo* (**Figures 2D and 2E**). The half-decay time of the cAMP transients was around 20 seconds, in contrast to the Ca^2+^ transients on the order of hundreds of milliseconds (**Figure 2F**).

**Figure 2.**
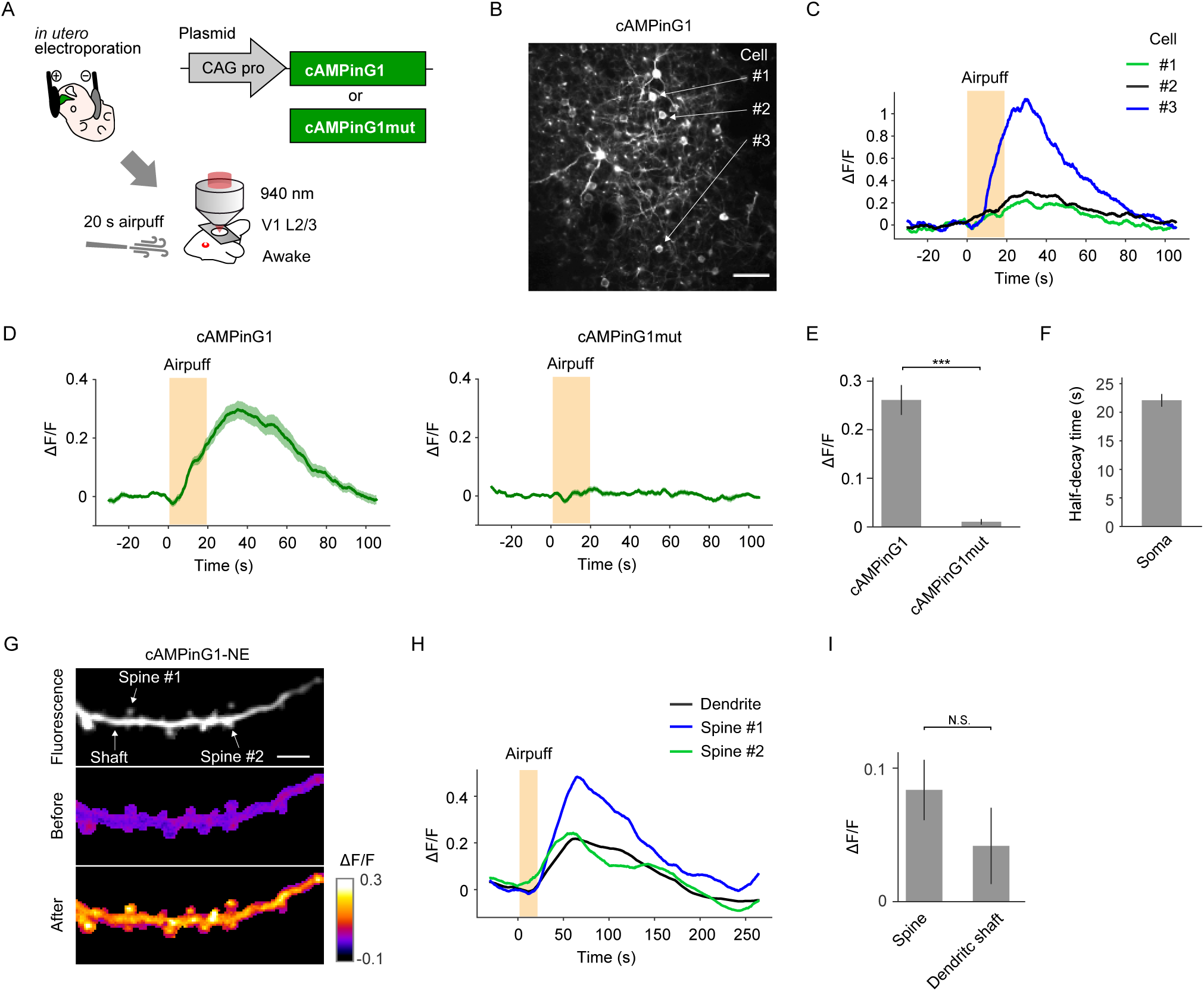
*In vivo* two-photon cAMPinG1 imaging in somata and dendritic spines. **(A)** Schematic of the experimental procedure of cAMPinG1 somatic imaging. cAMPinG1 or cAMPinG1mut was delivered to neurons in layer 2/3 (L2/3) of the mouse primary visual cortex (V1) by *in utero* electroporation. **(B)** A representative *in vivo* two-photon fluorescence image of cAMPinG1. Scale bar, 50 μm. **(C)** Single-trial cAMP traces of representative 3 cells. The orange box indicates the timing of the stimulus. **(D)** Averaged traces of somatic signals of cAMPinG1 (left) and cAMPinG1mut (right) in response to airpuff stimulation. n = 47 neurons in 4 mice (cAMPinG1), n = 39 neurons in 4 mice (cAMPinG1mut). **(E)** Averaged ΔF/F of cAMPinG1 and cAMPinG1mut in response to airpuff stimulation. n = 47 neurons in 4 mice (cAMPinG1), n = 39 neurons in 4 mice (cAMPinG1mut). Unpaired t-test. **(F)** Half-decay time of somatic cAMP transients in response to airpuff. n = 47 neurons in 4 mice. **(G)** A representative image of cAMPinG1 imaging in spines and their shaft. cAMPinG1 fluorescence (top), ΔF/F before (middle) and after (bottom) airpuff. Scale bar, 5 μm. **(H)** Representative traces of a dendritic shaft and two spines. The orange square indicates the timing of the stimulus. **(I)** Averaged ΔF/F of cAMPinG1 in dendritic shafts and spines. n = 56 spines, n = 11 shafts in 4 mice. Unpaired t-test. All shaded areas and error bars denote the SEM.

To further demonstrate the utility of cAMPinG1 indicators for cellular compartments, we expressed cAMPinG1-NE by *in utero* electroporation and imaged cAMP signals in dendritic spines and shafts *in vivo* under lightly anesthetized conditions (**Figure 2G**). We observed robust sensory-evoked cAMP transients in both dendritic spines and shafts (**Figures 2H and 2I**). These results show the feasibility of cAMPinG1 for *in vivo* imaging at both cellular and subcellular resolutions.

### Engineering and characterization of improved red Ca^2+^ indicator RCaMP3

Ca^2+^ is one of the most important intracellular signaling molecule that play a crucial role in regulating various physiological functions in neurons. However, the relationship between Ca^2+^ and cAMP remains poorly understood. To shed light on this relationship, it is essential to measure the dynamics of Ca^2+^ and cAMP simultaneously. For combinational use with the green cAMP indicator *in vivo*, we developed a new red Ca^2+^ indicator. Since the first circularly permuted red fluorescent protein (cpRFP)-type Ca^2+^ indicator, R-GECO1, was reported, a series of red Ca^2+^ indicators based on R-GECO1 have been developed (Dana et al., 2016; Inoue et al., 2015; Inoue et al., 2019; Ohkura et al., 2012; Wu et al., 2013; Zhao et al., 2011) (**Figure S4A**). We introduced several mutations from the existing Ca^2+^ indicators into jRGECO1a and termed this new hybrid design of the red calcium indicator RCaMP3 (**Figure 3A**).

**Figure 3.**
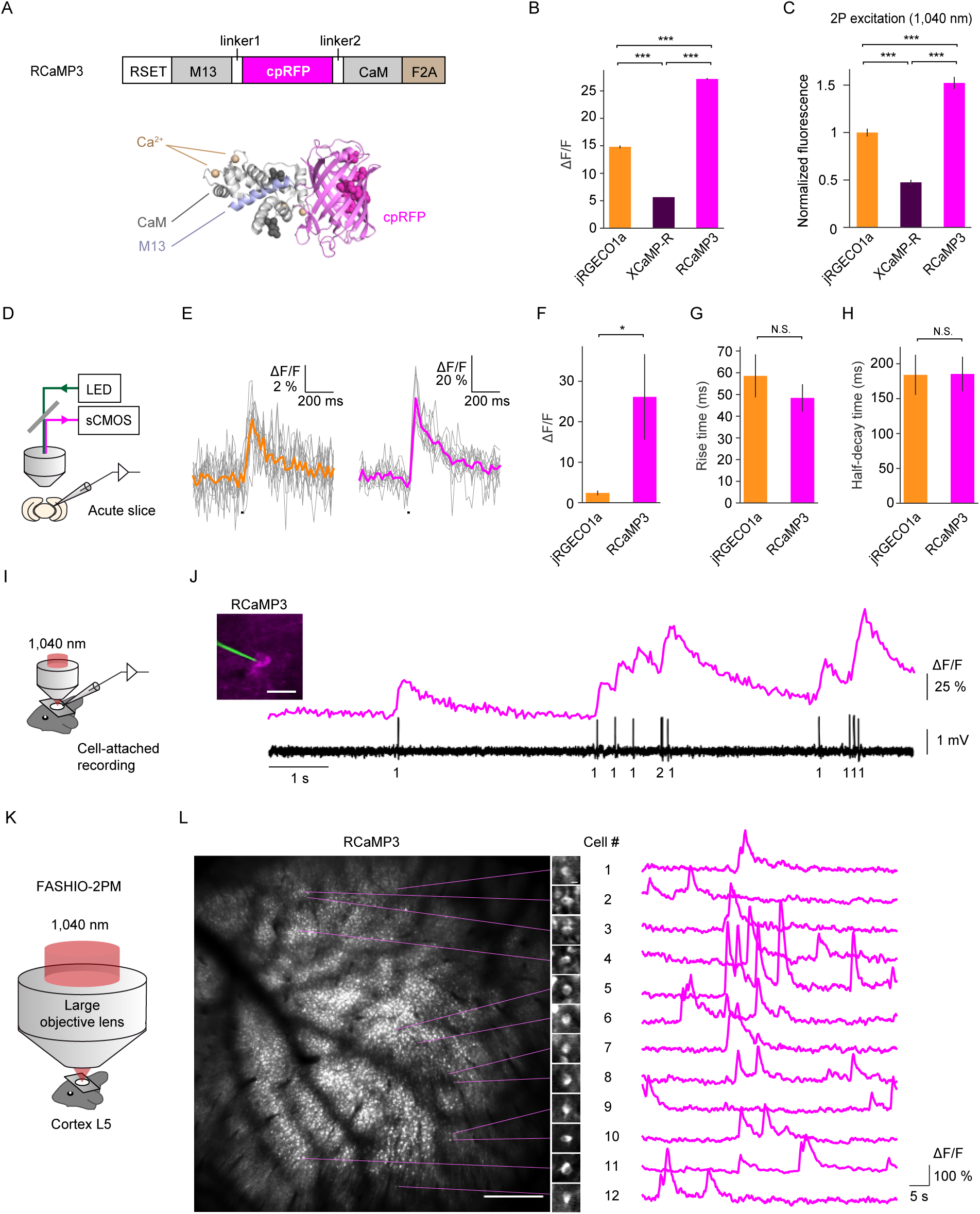
Engineering and characterization of RCaMP3. **(A)** Top: Primary structure of RCaMP3. The location of substitutions relative to R-GECO1 is indicated in R-GECO1 numbering. Bottom: Tertiary structures of R-GECO1 (PDB: 4I2Y) depicted as ribbon diagrams. Amino acids mutated in RCaMP3 are indicated in sphere shape. (B) ΔF/F of red Ca^2+^ indicators in HEK cell lysate. RCaMP3 had the largest ΔF/F. n = 4 wells (jRGECO1a), n = 4 wells (XCaMP-R), n = 4 wells (RCaMP3). Tukey’s post hoc test following one-way ANOVA. (C) Two-photon (1,040 nm) fluorescence intensities of red Ca^2+^ sensors in live HEK cells in the presence of ionomycin application. n = 360 (jRGECO1a), n =267 (XCaMP-R), n =376 (RCaMP3) cells. Tukey’s post hoc test following one-way ANOVA. (D) Schematic of the experimental procedure of Ca^2+^ imaging under a whole-cell patch-clamp configuration in acute brain slices. **(E)** Representative Ca^2+^ traces of jRGECO1a and RCaMP3 in response to a single action potential. Grey lines denote individual traces (10 trials), and colored thick lines denote average response. The black vertical lines indicate stimuli. **(F-H)** ΔF/F (F), rise time (G), and half-decay time (H) of jRGECO1a and RCaMP3 in response to a single action potential. n = 7 neurons (jRGECO1a), n = 6 neurons (RCaMP3). **(I)** Schematic of the experimental procedure of Ca^2+^ imaging under a cell-attached recording *in vivo*. **(J)** Representative trace of simultaneous measurement of RCaMP3 fluorescence and action potentials *in vivo*. The number of spikes for each event is indicated below the trace. The image shows a neuron expressing RCaMP3 (magenta) with the recording pipette (green). Scale bar, 20 μm. **(K)** Schematic of the experimental procedure of two-photon mesoscale Ca^2+^ imaging using fast-scanning high optical invariant two-photon microscopy (FASHIO-2PM). Cortical layer 5 (L5) neurons in the field-of-view (FOV, 3.0 × 3.0 mm^2^) were imaged by 1,040 nm excitation. **(L)** Left: A representative full FOV of FASHIO-2PM. Scale bar, 500 μm. Right: Magnified images and Ca^2+^ traces of representative 12 neurons. Scale bar, 10 μm. All error bars denote the SEM.

*In vitro* characterization revealed that RCaMP3 had a larger dynamic range and more blue-shifted excitation spectra compared to jRGECO1a and XCaMP-R, two of the best red calcium indicators for *in vivo* two-photon imaging (**Figure 3B; Figures S4B-S4F**). In addition, RCaMP3 exhibited a similar Ca^2+^ sensitivity and Hill coefficient to jRGECO1a (**Figures S4G-S4H**). Some of the most commercially available two-photon lasers are equipped with fixed around 1,040 nm for excitation of red fluorophores though the two-photon spectral peak of jRGECO1a and XCaMP-R is longer than 1,040 nm (Inoue et al., 2019). When excited at 1,040 nm, RCaMP3 showed a significant increase in fluorescence over jRGECO1a in the Ca^2+^-saturated state (**Figure 3C**). Therefore, the enhanced dynamic range and blue-shifted excitation spectrum of RCaMP3 make it particularly well-suited for two-photon imaging using the commonly available 1,040 nm lasers.

Next, we tested the performance of RCaMP3 in acute brain slices of L2/3 barrel cortex pyramidal neurons introduced by adeno-associated virus (AAV). Spike-induced calcium transients were assessed by one-photon imaging under a whole-cell patch-clamp configuration (**Figure 3D**). We found that RCaMP3 had larger responses to single action potentials than jRGECO1a in response, while their rise and decay kinetics were comparable (**Figures 3E-3H**). In addition, we performed loose-seal cell-attached electrical recording and two-photon Ca^2+^ imaging simultaneously *in vivo* (**Figure 3I**). RCaMP3 reliably detected Ca^2+^ transients preceded by single APs (**Figure 3J**). These results indicate that RCaMP3 is superior to the existing red Ca^2+^ indicators in detecting spike-induced calcium transients with single cell resolution.

We then tested the performance of RCaMP3 in *in vivo* two-photon mesoscale imaging using fast-scanning high optical invariant two-photon microscopy (FASHIO-2PM) (Ota et al., 2021). We performed RCaMP3 imaging in large field-of-view (3.0 × 3.0 mm^2^) including the primary somatosensory cortices by 1,040 nm excitation (**Figure 3K; Movie S1**). Somatic Ca^2+^ transients of several thousands of L5 neurons were simultaneously monitors with single-cell resolution (**Figure 3L**). In addition, somatic Ca^2+^ transients of L2/3 neurons and dendritic Ca^2+^ transients of L5 neurons could be monitored when imaged in L2/3 (**Figure S5**). These results demonstrate the high sensitivity of RCaMP3 which enables Ca^2+^ imaging large sets of deep cortical neurons.

### Dual-color imaging for Ca^2+^ and cAMP during forced running

Next, to image the Ca^2+^ and cAMP dynamics with single-cell resolution *in vivo*, we co-expressed RCaMP3 and cAMPinG1-ST in L2/3 neurons of the V1 by AAV injection (**Figure 4A**). Using a piezo objective scanner, we performed multiple z-plane imaging of head-fixed awake mice (**Figure 4B**). cAMPinG1-ST and RCaMP3 were excited with 940 nm and 1,040 nm, respectively. Consistent with the previous report, cAMPinG1-ST fluorescence was localized somata due to the soma-targeting signal RPL10 (Chen et al., 2020), which reduced contamination of neuropil fluorescence and enabled accurate tracking of individual cAMP changes. Here, we simultaneously visualized Ca^2+^ and cAMP signals of more than 400 neurons in L2/3 of a mouse (**Figures 4C and 4D; Movie S2**). During the imaging, we employed a forced running task, which was reported to increase neuromodulators, including noradrenaline, cAMP, and PKA activities in the cortex (Ma et al., 2018; Massengill et al., 2022; Wang et al., 2022). The majority of neurons showed an increase in cAMP signals during running (**Figures 4D and 4E**). This global cAMP increase was less cell-specific, possibly due to neuromodulators such as noradrenaline (Massengill et al., 2022; Reimer et al., 2016). Consistent with the previous report (Stringer et al., 2019), some parts of cells showed calcium transients during running, detected by RCaMP3 (**Figures 4D and 4E**). Interestingly, these motion-related cells had larger cAMP transients than the other non-motion-related cells (**Figures 4F-4H**). This additional cAMP elevation with Ca^2+^ responses may be attributed to Ca^2+^-dependent ACs. These results suggest that our multicolor imaging can detect Ca^2+^ and cAMP signals separately, and that cAMP signals can integrate information encoded in multiple upstream neuromodulators and Ca^2+^.

**Figure 4.**
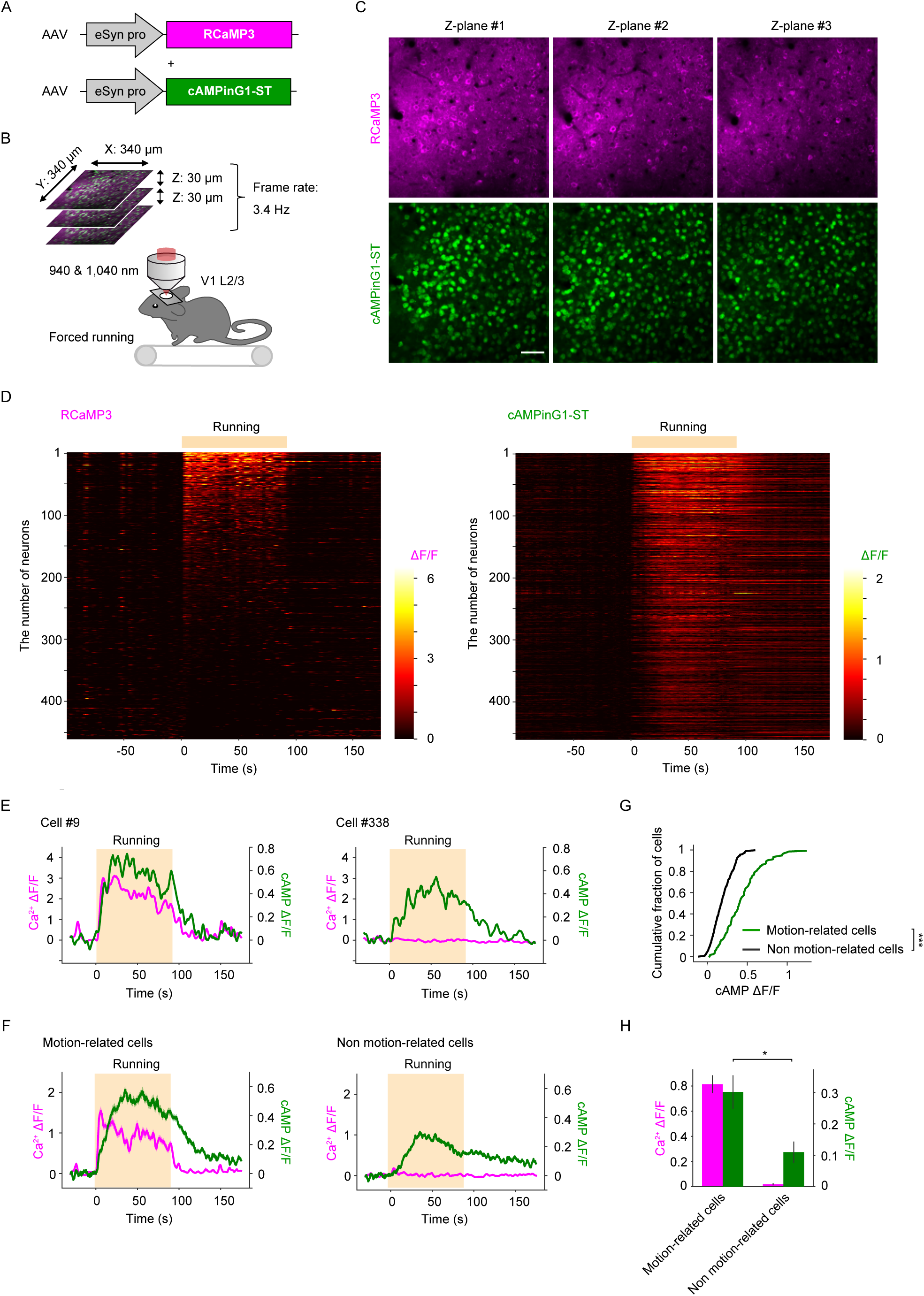
*In vivo* dual-color imaging for Ca^2+^ and cAMP during forced running. **(A)** Schematic of AAVs. AAVs encoding RCaMP3 and cAMPinG1-ST were co-injected into the L2/3 of the V1. **(B)** Schematic of the experimental procedure. Sequential excitation at 940 nm and 1,040 nm was used for dual-color imaging of cAMPinG1-ST and RCaMP3. Three optical planes spaced 30 µm apart were imaged at 3.4 Hz per plane using a piezo objective scanner. **(C)** Representative images of RCaMP3 and cAMPinG1-ST. Scale bar, 50 μm. **(D)** Single-trial traces of RCaMP3 and cAMPinG1-ST. Cells are sorted according to ΔF/F of RCaMP3 during running. n = 461 cells in 1 mouse. **(E)** Single-trial traces of RCaMP3 (magenta) and cAMPinG1-ST (green) of two representative cells. The orange box indicates the period of forced running. The cell number on the top corresponds to the number in (D). **(F)** Averaged fluorescence transients of RCaMP3 (magenta) and cAMPinG1-ST (green). n = 137 cells in 1 mouse (left), n = 324 cells in 1 mouse (right). **(G)** Cumulative plot of mean cAMP ΔF/F of motion-related (green) and non-related (black) cells during forced running. n = 137 cells in 1 mouse (green), n = 324 cells in 1 mouse (black). Kolmogorov–Smirnov test. **(H)** Averaged ΔF/F of RCaMP3 (magenta) and cAMPinG1-ST (green) during forced running. n = 3 mice. Paired t-test. All shaded areas and error bars denote the SEM.

To demonstrate the application of cAMPinG1 and RCaMP3 for other cell types than neurons, we expressed RCaMP3 and cAMPinG1-NE in astrocytes in L2/3 of the V1 by viral delivery of them under the control of the GFAP promoter, which introduced specific expression in astrocytes (Lee et al., 2006). Again, forced running-induced Ca^2+^ increase followed by a cAMP increase (**Figure S6**).

### Dual-color imaging for Ca^2+^ and cAMP during visual stimulation

To further investigate the relationship between Ca^2+^ and cAMP *in vivo*, a drifting grating stimulus of 8 directions was applied to induce cell-specific Ca^2+^ transients in L2/3 neurons of the V1 (**Figure 5A**). *In vivo* two-photon imaging revealed that RCaMP3 showed direction-selective Ca^2+^ transients in response to 4 seconds of drifting gratings, consistent with previous studies (Chen et al., 2013; Dana et al., 2016; Sakamoto et al., 2022a) **(Figures 5B-5D; Figure S7**). Interestingly, cAMPinG1-ST also showed direction-selective cAMP transients preceded by Ca^2+^ transients, which were selective to the same direction **(Figures 5B-5D; Figure S7**). These data suggest that as well as forced running, visual stimuli also induced cAMP increase preceded by Ca^2+^ transients.

**Figure 5.**
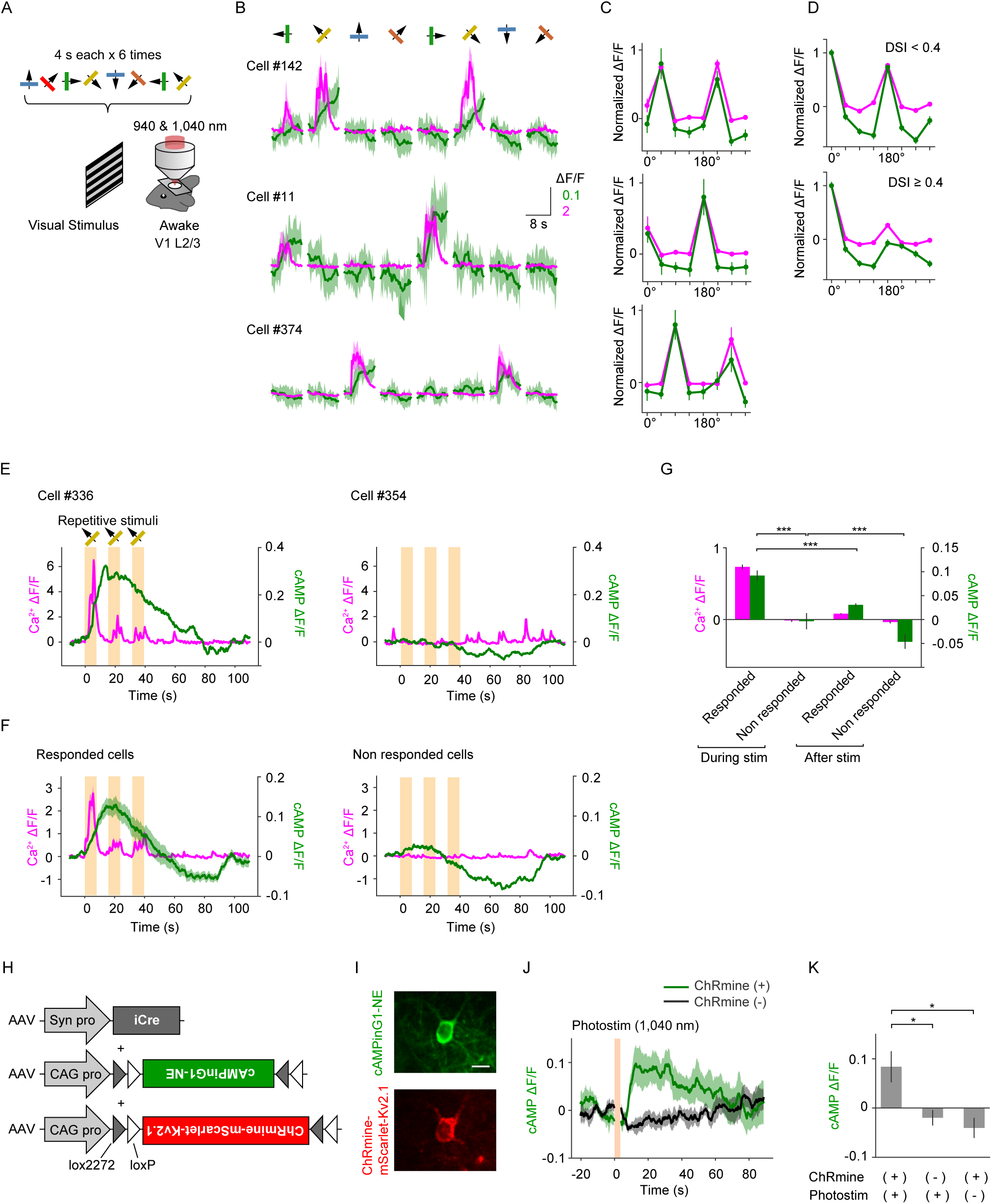
*In vivo* dual-color imaging for Ca^2+^ and cAMP during visual stimulation. **(A)** Schematic of the experimental procedure. Moving gratings of 8 directions were used to induce cell-specific Ca^2+^ transients in L2/3 neurons of the V1. **(B)** Averaged fluorescence transients of RCaMP3 (magenta) and cAMPinG1-ST (green) of 3 representative cells. **(C)** Direction-selective visual responses of RCaMP3 (magenta) and cAMPinG1-ST (green) of the 3 representative cells in (B). **(D)** Averaged direction-selective visual responses of RCaMP3 (magenta) and cAMPinG1-ST (green). Top: neurons showing a direction selectivity index (DSI) < 0.4 in Ca^2+^ response. n = 94 cells in 3 mice. Bottom: neurons showing a direction selectivity index (DSI) ≥ 0.4 in Ca^2+^ response. n = 101 cells in 3 mice. **(E)** Single-trial traces of RCaMP3 (magenta) and cAMPinG1-ST (green) of 2 representative cells. The orange box indicates the period of visual stimuli. **(F)** Averaged fluorescence transients of RCaMP3 (magenta) and cAMPinG1-ST (green). n = 53 cells in 1 mouse (left), n = 408 cells in 1 mouse (right). **(G)** Averaged ΔF/F of RCaMP3 (magenta) and cAMPinG1-ST (green) during and after the visual stimuli. n = 3 mice. Paired t-test. **(H)** Schematic of AAVs for sparse expression of cAMPinG1-NE and soma-targeted ChRmine. **(I)** Representative fluorescence images of cAMPinG1-NE and ChRmine-mScarlet-Kv2.1. Scale bar, 10 μm. **(J)** Averaged fluorescence transients of cAMPinG1-NE in response to 1,040 nm photostimulation. n = 11 neurons in 3 mice (ChRmine (+), green), n = 11 neurons in 3 mice (ChRmine (-), black). **(K)** Averaged ΔF/F of cAMPinG1-NE in response to 1,040 nm photostimulation. n = 11 neurons in 3 mice (ChRmine (+), photostim (+)), n = 11 neurons in 3 mice (ChRmine (-), photostim (+)), n = 9 neurons in 3 mice (ChRmine (+), photostim (-)). Tukey’s post hoc test following one-way ANOVA. All shaded areas and error bars denote the SEM.

To explore cAMP changes in response to visual stimuli in more detail, we conducted an experiment with repetitive moving grating of the same direction (8 seconds, 3 times). Neurons responding to the direction in Ca^2+^ levels exhibited cAMP increase, as observed above (**Figures 5E-5G**). Surprisingly, we observed that cAMPinG1-ST fluorescence began to decrease in majority of Ca^2+^-responsive and unresponsive cells in the middle of visual stimuli. This global cAMP decrease, lasted beyond the end of stimulus and lowered cAMPinG1 fluorescence than the baseline observed before the stimulation (**Figures 5E-5G**). This phenomenon was not observed during forced running, potentially due to differences in the neuromodulators, GPCRs and PDEs activated by these physical stimuli. Overall, dual-color Ca^2+^ and cAMP imaging during visual stimuli visualized cAMP levels modulated bidirectionally by multiple upstream *in vivo*.

### Action potentials are sufficient to induce somatic cAMP elevation

To determine whether the calcium-related cAMP increase detected above depended on Ca^2+^ influx induced by action potentials, we applied single-cell optogenetic stimulation and cAMP imaging *in vivo*. To achieve spatially precise optical stimulation to the soma of a neuron, we sparsely expressed cAMPinG1-NE and soma-targeted ChRmine in L2/3 neurons of the V1 by utilizing Cre/loxP recombination system (Druckmann et al., 2013; Marshel et al., 2019) (**Figures 5H and 5I**). Since ChRmine is a red-shifted cation-conducting channelrhodopsin that is selective to monovalent cations, its use allows for ruling out the effect of Ca^2+^ influx through the rhodopsin on Ca^2+^-dependent cAMP changes (Kishi et al., 2022). We observed a robust increase in cAMP levels in response to two-photon single-cell stimulation at 1,040 nm (**Figure 5J**). This cAMP increase is necessary for both the expression of ChRmine and photostimulation, meaning that the spike-induced cAMP increase can be mediated by voltage-dependent Ca^2+^ channels and Ca^2+^-dependent ACs (**Figure 5K**). These data indicate that action potentials are sufficient to induce an increase in somatic cAMP level through the Ca^2+^ signaling pathway.

### Dual-color fiber photometry for Ca^2+^ and cAMP

Having demonstrated the superior performance of cAMPinG1 and RCaMP3 by *in vivo* two-photon imaging, we next evaluated their potential use in single photon fiber photometry in deep brain areas. We injected AAVs encoding RCaMP3 and cAMPinG1-NE into the dorsal striatum (dStr) (**Figure 6A**). Since the neural activity of the dStr is associated with body movements (Parker et al., 2018), we performed the dual-color fiber photometry to measure Ca^2+^ and cAMP levels during a forced running task. For dual-color imaging, we employed different excitation wavelengths: 560 nm for RCaMP3 imaging and 405 nm and 470 nm for cAMPinG1 ratiometric imaging. In line with previous studies (Massengill et al., 2022; Wang et al., 2022), we observed an increase in cAMP levels during the running period following the increase in Ca^2+^ levels (**Figures 6B-6D**). Surprisingly, cAMP levels decreased after the running period and remained low for tens of seconds. In addition, we found that cAMPinG1mut-NE did not show clear fluorescent change during and after the running task (**Figures 6B-6D**). These data indicate that dual-color fiber photometry using RCaMP3 and cAMPinG1-NE can detect bulk Ca^2+^ increase and bidirectional cAMP changes in deep brain regions.

**Figure 6.**
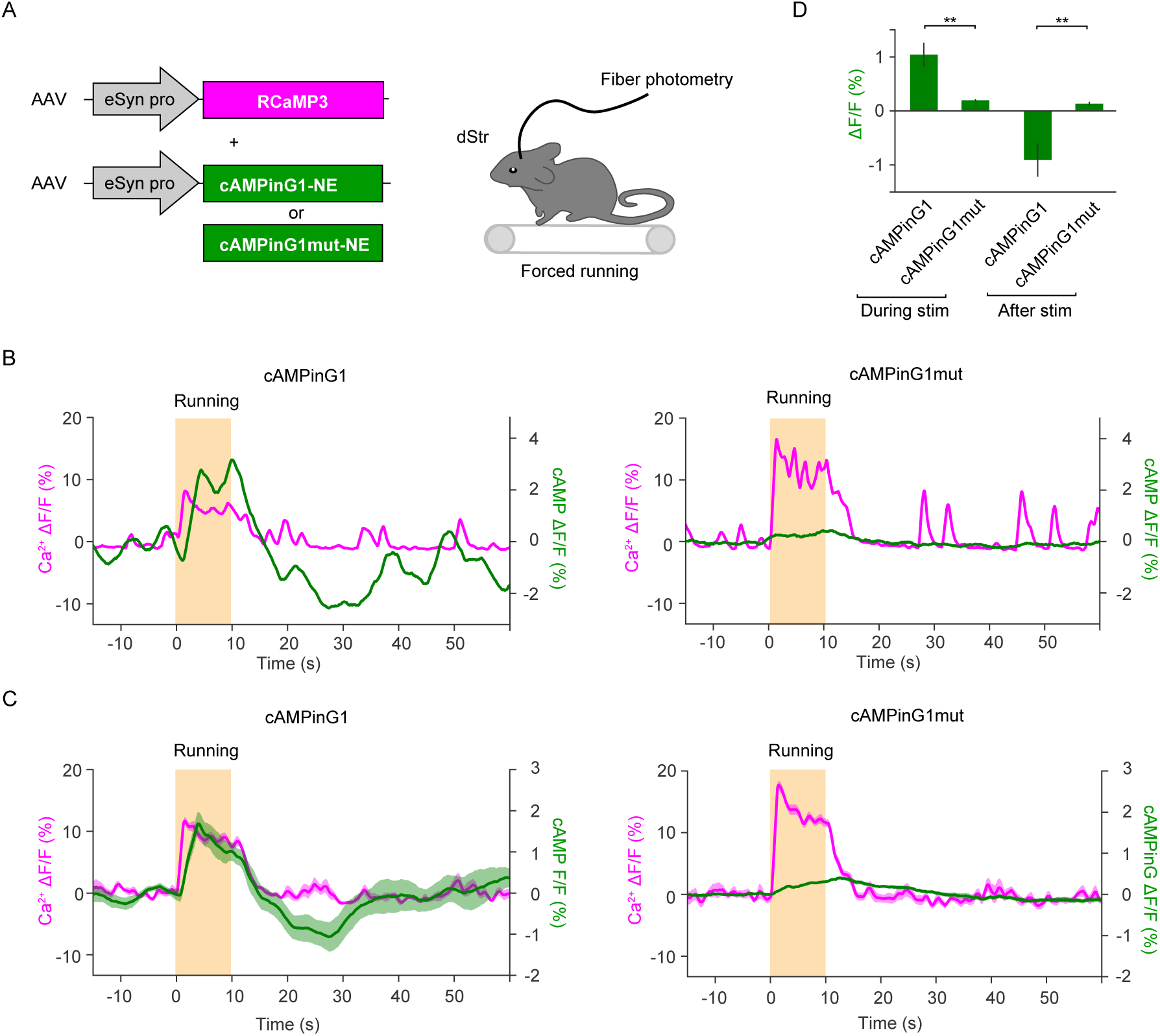
Dual-color fiber photometry for Ca^2+^ and cAMP. **(A)** Schematic of the experimental procedure. Dual-color fiber photometry was performed in the dorsal striatum (dStr) during a forced running task. **(B)** Representative single-trial traces of cAMPinG1-NE (green, left), cAMPinG1mut-NE (green, right), and RCaMP3 (magenta) signals. The orange box indicates the period of forced running. **(C)** Averaged fluorescence traces of cAMPinG1-NE (green, left), cAMPinG1mut-NE (green, right), and RCaMP3 (magenta) signals. n = 27 trials in 3 mice (left), n = 27 trials in 3 mice (right). **(D)** Averaged ΔF/F of cAMPinG1-NE and cAMPinG1mut-NE during and after the stimulation. n = 27 trials in 3 mice (cAMPinG1-NE), n = 27 trials in 3 mice (cAMPinG1mut-NE). Unpaired t-test. All shaded areas and error bars denote the SEM.

### Single-cell, single-timepoint cAMPinG1 imaging for GPCR biology and drug screening

Finally, to demonstrate the utility of cAMPinG1 for cAMP quantification in GPCR biology and drug screening, we generated a stable cell line of HEK293T cells that express cAMPinG1. Single-cell cloning of the stable cell lines resulted in the homogenous expression levels of cAMPinG1 among all cells. The proliferation rate of the cAMPinG1 stable cell line was comparable with the original HEK293T cell line, indicating the undetectable toxicity of cAMPinG1 expression (**Figures S8A and S8B**). As previously shown (**Figures 1C and 1I-1K**), cAMPinG1 is more suitable for ratiometric imaging compared to other cpGFP-based cAMP indicators. Therefore, we performed ratiometric imaging using the cAMPinG1 stable cell line. After applying forskolin, we found that the 488 ex / 405 ex ratio in each cell was higher in forskolin-stimulated cells than in non-stimulated cells (**Figure S8C**). This result shows that ratiometric cAMPinG1 imaging can be utilized to quantify cAMP levels without the need for time-lapse imaging, simply comparing the 488 ex / 405 ex ratio at a specific point in time before and after drug administration with cellular resolution. We referred to this method as “single-timepoint cAMPinG1 imaging.”

To demonstrate the utility of single-timepoint cAMPinG1 imaging for GPCR biology, we transiently expressed a dopamine receptor D1 (DRD1), a Gs-coupling GPCR known for its constitutive activity in the cAMP pathway, in the cAMPinG1 stable cell line (**Figure 7A**). We fused a self-cleaving 2A peptide (P2A) and RFP to DRD1 to monitor the expression level of DRD1 in each cell. We also used a tetracycline-inducible expression system to minimize the effect of GPCR expression on cell proliferation or toxicity. Single-cell, single-timepoint ratiometric imaging with cAMPinG1 revealed a positive correlation between DRD1 expression and cAMP level (Wang et al., 2020) (**Figures 7A and 7B**). Consistent with previous reports (Lin et al., 2020; Wang et al., 2020), we also observed high cAMP levels due to the constitutive activity of the GPCR GPR52 (**Figure 7E**). These results suggest that single-timepoint cAMPinG1 imaging can be available to compare the constitutive activity of cAMP-related GPCRs. In addition to measuring constitutive activities, single-timepoint cAMPinG1 imaging also allowed quantifying the agonist activity of a Gi-coupling dopamine receptor D2 (DRD2) as well as that of DRD1 with cellular resolution (**Figures 7C and 7D, 7F**). Furthermore, to demonstrate the utility of cAMPinG1 imaging for detecting the inverse agonist activity, which reduces the constitutive activity of GPCRs, we used a Gs-coupling serotonin receptor HTR6. The activity of inverse agonist clozapine was detected as well as the agonist activity of serotonin (**Figure 7F**). These data demonstrate the utility of single-timepoint cAMPinG1 imaging for GPCR biology and pharmacology.

**Figure 7.**
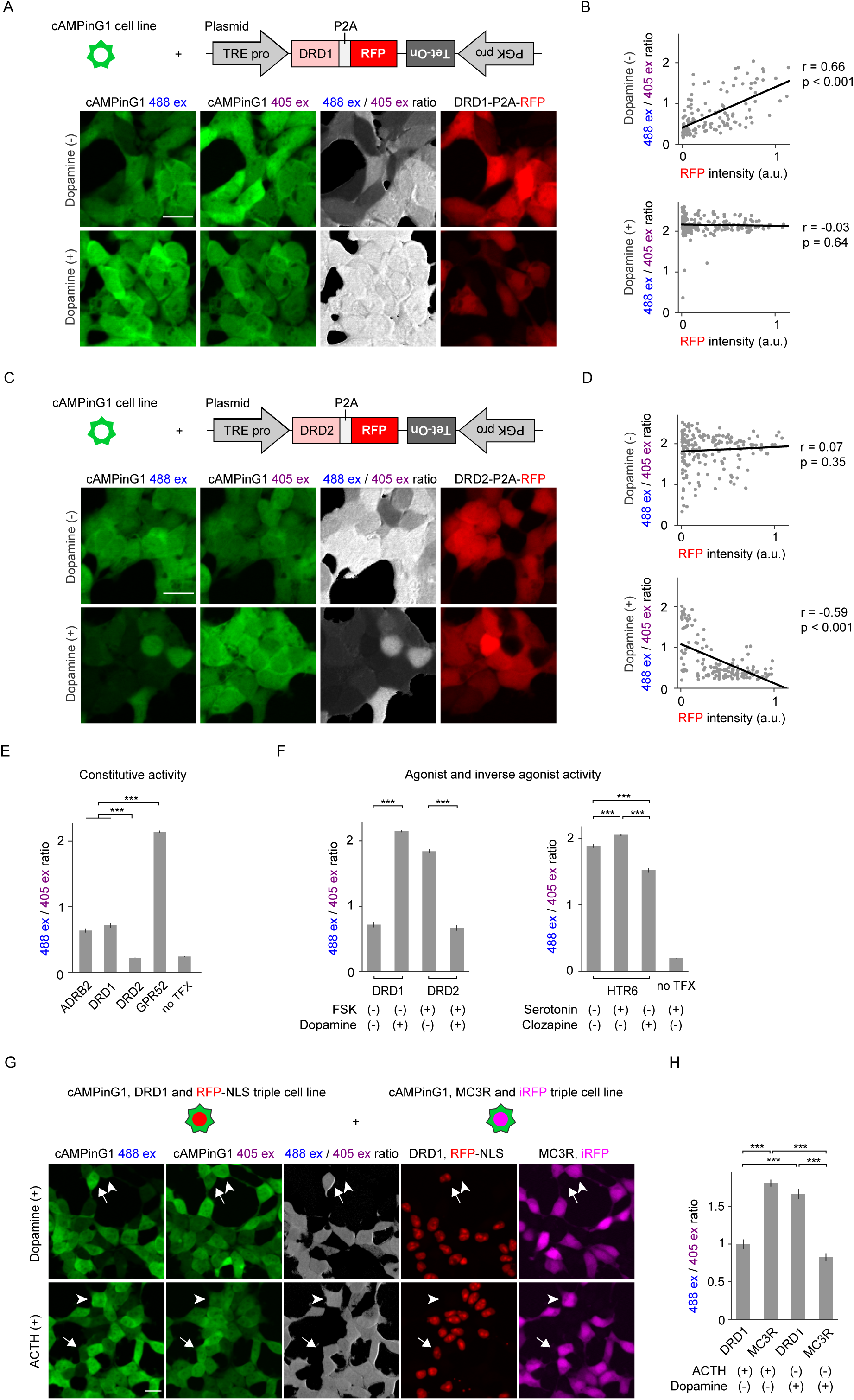
Single-cell, single-timepoint cAMPinG1 imaging for GPCR biology and drug screening. **(A)** Schematic of the expression system (top). The plasmid encoding pTRE-DRD1-P2A-mCherry-reverse-PGK-TetOn was transfected into a cAMPinG1 stable cell line. Doxycycline was added 3 hours before the imaging to induce expression of DRD1-P2A-mCherry. Representative images of cAMPinG1 stable cell line expressing DRD1-P2A-RFP in the absence (middle) or presence (bottom) of 1,000 nM dopamine were taken by alternating 405/488/561 nm lasers excitation. Scale bar, 20 μm. **(B)** Correlation between blue ex/violet ex ratio of cAMPinG1 and DRD1-P2A-RFP expression level. Individual dots indicate single cells. n = 141 (dopamine (-), top) and 190 (dopamine (+), bottom) cells. Pearson correlation coefficient in linear regression, r = 0.66; *p* < 0.001 (top) and r = -0.03; *p* = 0.64 (bottom). **(C)** Schematic of the expression system (top). Representative images of a cAMPinG1 cell line expressing DRD2-P2A-RFP in the absence (middle) or presence (bottom) of 1,000 nM dopamine were taken by alternating 405/488/561 nm lasers excitation. 0.5 μM forskolin was applied in both conditions. Scale bar, 20 μm. **(D)** Correlation between blue ex/violet ex ratio of cAMPinG1 and DRD2-P2A-RFP expression level. Individual dots indicate single cells. n = 180 (dopamine (-), top) and 154 (dopamine (+), bottom) HEK293T cells. Pearson correlation coefficient in linear regression, r = 0.07; *p* = 0.35 (top) and r = -0.59; *p* < 0.001 (bottom). **(E)** The blue ex/violet ex ratio of cAMPinG1 cells transiently expressing GPCRs-P2A-RFP in the absence of ligands, indicating he constitutive activity of each GPCR. n = 158 (ARDB2), 141 (DRD1), 151 (DRD2), 178 (GPR52), 126 (no TFX) cells. Tukey’s post hoc test following one-way ANOVA. **(F)** Blue ex/violet ex ratio of cAMPinG1 cell line expressing GPCRs-P2A-RFP with or without agonists. Note that clozapine is known to act as an inverse agonist against HTR6. n = 141 (DRD1), 190 (Dopamine + DRD1), 180 (FSK + DRD2), 154 (Dopamine + FSK + DRD2) cells (left). n = 159 (HTR6), 151 (5HT + HTR6), 135 (Clozapine + HTR6), 204 (5HT + no TFX) cells (right). Tukey’s post hoc test following one-way ANOVA. **(G)** Schematic of the expression system (top). Representative images of a mixture of cell lines stably expressing cAMPinG1, DRD1 and RFP, and cell lines stably expressing cAMPinG1, MC3R and iRFP in the presence of 100 nM dopamine (middle) or 1,000nM ACTH (bottom). Representative cells expressing DRD1/RFP or MC3R/iRFP were indicated by arrows or arrowheads, respectively. Scale bar, 20 μm. **(H)** The blue ex/violet ex ratio of each cell line in the presence of dopamine or ACTH. Both cell line showed ligand-specific cAMP elevation. n = 95 (ACTH + DRD1), 87 (ACTH + MC3R), 72 (dopamine + DRD1), 161 (dopamine + MC3R) cells. Tukey’s post hoc test following one-way ANOVA. All error bars denote the SEM.

To further demonstrate the utility of single-timepoint imaging as a method for drug screening, we established triple stable cell lines expressing cAMPinG1, a GPCR, and a marker fluorescent protein. We used DRD1 and melanocortin 3 receptor (MC3R) as representative Gs-coupling GPCRs. We mixed DRD1-RFP expressing cells and MC3R-iRFP expressing cells to image the responsiveness of each GPCR simultaneously (**Figure 7G**). Single-timepoint cAMPinG1 imaging revealed that DRD1 and MC3R specifically responded to their respective agonist (dopamine and ACTH, respectively) (**Figure 7H**). These results suggest that single-timepoint cAMPinG1 imaging has the potential for multiplex high-throughput drug screenings, allowing for the simultaneous evaluation of multiple GPCRs and exclusion of compound candidates that increase cAMP levels in multiple GPCR cell lines and produce non-specific responses.

## DISCUSSION

### Engineering of cAMPinG1 and RCaMP3

In this study, we report the engineering of cAMPinG1, an ultrasensitive cAMP indicator with high affinity for cAMP and a large dynamic range. The sensitivity was enough to quantify cAMP transient in both somata and dendritic spines in the mouse cortex. In addition, by combining cAMPinG1 with RCaMP3, we visualized the population dynamics of cAMP and Ca^2+^ simultaneously in hundreds of cells with cellular resolution *in vivo*. We also conducted proof-of-concept experiments of cAMPinG1 imaging for GPCR biology and drug screening.

Although several cAMP sensors have been developed recently, each had some drawbacks for *in vivo* imaging (Massengill et al., 2022; Wang et al., 2022). Due to its low cAMP affinity, G-Flamp1 limited the number of cells that could be imaged and quantified cAMP dynamics in the mouse cortex. cAMPFIREs were applied for two-photon fluorescence lifetime imaging microscopy (FLIM), which made fast imaging or combined use with other sensors, such as Ca^2+^ indicators, generally challenging due to the required optical settings. In addition, the full-length cAMPFIREs cDNA (approximately 4.5 kb) is too large to be packaged into a AAV vector, limiting on the methods for *in vivo* delivery. Our new indicators (about 1.7 kb) with high sensitivity addressed these issues. To the best of our knowledge, this is the first study to visualize population dynamics of Ca^2+^ and cAMP simultaneously in hundreds of cells in the mouse cortex, demonstrating a robust positive correlation between Ca^2+^ and cAMP.

Red Ca^2+^ indicators are indispensable for multicolor imaging with a green indicator to reveal the interaction between the target molecule of the green indicator and neural activity or Ca^2+^ signaling. Despite tremendous efforts to improve the sensitivity of red calcium indicators (Dana et al., 2016; Fenno et al., 2020; Inoue et al., 2015; Inoue et al., 2019; Ohkura et al., 2012; Wu et al., 2013; Zhao et al., 2011), the application of red Ca^2+^ indicators is still limited due to their sensitivity *in vivo*. In this study, we developed a highly sensitive RCaMP3, which expands the application of red Ca^2+^ indicators for two-photon mesoscale imaging. Moreover, multicolor imaging of RCaMP3 and cAMPinG1-ST revealed the interaction of Ca^2+^ and cAMP at the population level. Therefore, RCaMP3 will contribute to further multicolor imaging, especially with recently developed green indicators (Duffet et al., 2022; Fenno et al., 2020; Ino et al., 2022; Unger et al., 2020).

### Population coding in Ca^2+^ and cAMP *in vivo*

*In vivo* imaging experiments showed bidirectional cAMP changes in response to various physiological stimuli. In the cortex, it is reported that noradrenaline is secreted in response to forced running or aversive stimuli such as airpuffs and activates Gs-coupling β1 adrenoceptors (Massengill et al., 2022; Oe et al., 2020; Reimer et al., 2016; Wang et al., 2022), which can explain the global cAMP increase observed in this study. On the contrary, there are multiple possibilities to explain the mechanisms of the global cAMP decrease observed during and after stimulations in the cortex and dorsal striatum, including some Gi-coupling GPCRs, such as GABA_B_ receptors, acetylcholine receptors, and adrenergic receptors, or PDEs activated by Ca^2+^ or kinases (Omori and Kotera, 2007). Furthermore, the detection of downward cAMP change indicates the baseline cAMPinG1 signal before stimulation reflects basal cAMP level, which is the sum of constitutive activities of expressed GPCRs and agonist activities of the extracellular ligands existing originally in the awake conditions. Thus, the cAMP affinity of cAMPinG1, on the order of several hundreds of nanomolar, is suitable for baseline cAMP level and bidirectional cAMP change *in vivo*. In addition, our results showed that the kinetics of cAMP varied between the experiments, possibly due to the difference in serving GPCRs or expression properties of phosphodiesterase families in individual cell types or brain regions. Overall, the sensitivity of cAMPinG1 is suitable for visualizing bidirectional change and kinetics of cAMP in various cell types.

Two-photon dual-color imaging for Ca^2+^ and cAMP revealed a strong correlation between cell-specific cAMP increases and Ca^2+^ transients. Optogenetic experiments also showed that action potentials are sufficient to induce somatic cAMP transients *in vivo*. The spike-induced cAMP increase can be mediated by voltage-dependent Ca^2+^ channels and Ca^2+^-dependent ACs. Thus, our results provide the first evidence that cAMP can encode specific information, such as direction selectivity in vision or locomotion encoded in action potentials and Ca^2+^ signaling (**Figure S9A**). This cell-specific cAMP elevation cooperated or competed with global cAMP increase or decrease, respectively, leading to the formation of population patterns of cAMP. These forms of population coding in cAMP highlight the importance of Ca^2+^ signaling as the upstream regulator of cAMP and PKA signaling, which is not fully understood due to technological limitations (Ma et al., 2022; Reimer et al., 2016; Tang and Yasuda, 2017). Notably, cAMP transients last for tens of seconds, much longer than the hundreds of milliseconds of Ca^2+^ transients. Therefore, information encoded in Ca^2+^ and GPCR signaling is integrated and stored for a longer period through cAMP transients (**Figure S9B**).

### cAMPinG1 imaging for GPCR biology and drug screening

The human genome code about non-olfactory 300 GPCRs that are expressed throughout the body, and even small parts of GPCRs are related to one-third of all approved drugs, indicating the importance of studying GPCR biology and conducting drug screening targeting these receptors. Quantification of the constitutive activity of GPCRs is important because it has various roles *in vivo* (Iino et al., 2020; Wang et al., 2020). Moreover, the downstream pathway of orphan GPCRs, whose endogenous ligands are unknown, has been assessed by constitutive activity (Kroeze et al., 2015; Schihada et al., 2021). However, current technologies for measuring the constitutive activity of GPCRs have limitations, including the inability to cancel out the effects of GPCRs expression levels and cell proliferation rate. To overcome these limitations, we utilized a tetracycline-inducible expression system to minimize cell toxicity due to the expression of some GPCRs and single-cell, single-timepoint cAMPinG1 imaging to simultaneously assess cAMP levels and GPCRs-P2A-RFP expression levels. The single-cell analysis will also contribute to multiplex high-throughput screening, in which several GPCRs can be screened simultaneously combined with barcoding fluorescent proteins, helping to minimize false positives (Yang et al., 2021). Furthermore, our indicators can be used for inverse agonist screening due to their high sensitivity. Taken together, our brief and robust cAMP quantification technique has the strong potential to be valuable in GPCR biology and drug screening.

### Limitations of the study

Our results identified bidirectional cAMP change in the cortex and striatum, suggesting that Ca^2+^ and noradrenaline signaling may be upstream of cAMP. However, the functional roles of these cAMP changes in the brain network are also unclear because cAMP has multiple downstream pathways, including PKA and cAMP-dependent channels. Further investigation is necessary to fully understand these processes. It will also be important to determine the cAMP dynamics in different cell types in other brain areas or body parts because expression patterns of GPCRs, PDEs, and ACs vary among cell types. Overall, our precise *in vivo* imaging of Ca^2+^ and cAMP imaging will contribute to a variety of scientific fields.

Our study quantified the constitutive activity of several representative Gs-coupling GPCRs. It will be necessary to determine the cAMP-mediated downstream pathways of GPCRs by cAMP indicators because there are some discrepancies between the results of receptor-G protein ‘couplome’ obtained from different assays (Hauser et al., 2022). In addition, although GPCR signaling has several downstream pathways, such as IP3 and β-arrestin, our assay can only detect cAMP-related pathways. In the future, it will be important to develop more sensitive fluorescence indicators of these downstream molecules for GPCR biology and drug screening.

## METHODS

### KEY RESOURCES TABLE

| REAGENT or RESOURCE | SOURCE | IDENTIFIER |
| --- | --- | --- |
| Bacterial and virus strains |  |  |
| AAV2/1-eSyn-cAMPinG1-NE | This paper | N/A |
| AAV2/1-eSyn-cAMPinG1mut-NE | This paper | N/A |
| AAV2/1-eSyn-cAMPinG1-ST | This paper | N/A |
| AAV2/1-GFAP-cAMPinG1-NE | This paper | N/A |
| AAV2/1-CAG-DIO-cAMPinG1-NE | This paper | N/A |
| AAV2/1-eSyn-jRGECO1a | Dana et al., 2016 | N/A |
| AAV2/1-eSyn-RCaMP3 | This paper | N/A |
| AAV2/1-GFAP-RCaMP3 | This paper | N/A |
| AAV2/1-CAG-DIO-RCaMP3 | This paper | N/A |
| AAV2/1-hSyn-iCre | Druckmann et al., 2014 | N/A |
| AAVPHP.eB-pCaMKII-Cre | This paper | N/A |
| AAV2/1-CAG-DIO-ChRmine-mScarlet-Kv2.1 | Marshall et al., 2019 | N/A |
| LV-CAG-cAMPinG1 | This paper | N/A |
| LV-TRE3G-DRD1-cHS4-EF1 α-TetOn3G-P2A-mCherry-NLSx3 | This paper | N/A |
| LV-TRE3G-MC3R-cHS4-EF1 α-TetOn3G-P2A-iRFP670 | This paper | N/A |
| Chemicals, peptides, and recombinant proteins |  |  |
| cAMPinG1 | This paper | GenBank: xxxxxxxx |
| RCaMP3 | This paper | GenBank: xxxxxxxx |
| Cyclic adenosine monophosphate (cAMP) | Tokyo Chemical Industry | Cat# A2381 |
| Cyclic guanosine monophosphate (cGMP) | Abcam | Cat# ab120805 |
| Forskolin | Nacalai | Cat# 16384-84 |
| Forskolin | Wako | Cat# 067-02191 |
| IBMX | Tocris | Cat# 3758 |
| Dopamine Hydrochloride | Nacalai | Cat# 14212-71 |
| Clozapine | Cayman | Cat# 12059 |
| Adrenocorticotrophic hormone (ACTH) | AdooQ | Cat# AP3295 |
| Ionomycin (calcium salt) | Cayman Chemical | Cat# 11932 |
| Critical commercial assays |  |  |
| Calcium Calibration Buffer Kit#1 | Thermo Fisher Scientific | Cat# C3008MP |
| SYBR qPCR: SYBR Premix Ex Taq (Tli RNase H Plus) | Clontech | Cat# RR420 |
| NucleoBond Xtra Midi EF | MACHEREY-NAGEL | Cat# 740420 |
| Experimental models: cell lines |  |  |
| HEK293T | ATCC | Cat# CRL-11268 |
| DH5 $\alpha$ | TOYOBO | Cat# DNA-903 |
| DH10B | Invitrogen | Cat# 18297010 |
| Stbl3 | Invitrogen | Cat# C737303 |
| cAMPinG1 stable cell line | This paper | N/A |
| cAMPinG1, DRD1 and mCherry-NLS stable cell line | This paper | N/A |
| cAMPinG1, MC3R and iRFP670 stable cell line | This paper | N/A |
| Experimental models: organisms/strains |  |  |
| Mouse: ICR | Japan SLC | N/A |
| Mouse: C57BL/6N | Japan SLC | N/A |
| Recombinant DNA |  |  |
| pBAD-cAMPinG1 | This paper | N/A |
| pCAG-cAMPinG1 | This paper | RIKEN BRC: RDBxxxxx |
| pCAG-cAMPinG1mut | This paper | N/A |
| pCAG-cAMPinG1-NE | This paper | N/A |
| pCAG-cAMPinG1-ST | This paper | N/A |
| pCAG-Flamindo2 | Odaka et al., 2014 | N/A |
| pCAG-gCarvi | Kawata et al., 2022 | N/A |
| pCAG-G-Flamp1 | Wang et al., 2022 | N/A |
| pCAG-jRGECO1a | Dana et al., 2016 | N/A |
| pCAG-XCaMP-R | Inoue et al., 2019 | N/A |
| pCAG-RCaMP3 | This paper | N/A |
| pAAV-eSyn-cAMPinG1mut-NE | This paper | N/A |
| pAAV-eSyn-cAMPinG1-NE | This paper | RIKEN BRC: RDBxxxxx |
| pAAV-eSyn-cAMPinG1-ST | This paper | RIKEN BRC: RDBxxxxx |
| pAAV-CAG-DIO-cAMPinG1-NE | This paper | N/A |
| pAAV-gfaABC1D-cAMPinG1-NE | This paper | N/A |
| pAAV-eSyn-jRGECO1a | Dana et al., 2016 | N/A |
| pAAV-eSyn-RCaMP3 | This paper | RIKEN BRC: RDBxxxxx |
| pAAV-GFAP-RCaMP3 | This paper | N/A |
| pAAV-CAG-DIO-RCaMP3 | This paper | N/A |
| pAAV-hSyn-iCre | Druckmann et al., 2014 | N/A |
| pAAV-CaMKII-Cre | This paper | N/A |
| pAAV-CAG-DIO-ChRmine-mScarlet-Kv2.1 | Marshall et al., 2019 | N/A |
| pTRE-DRD1-P2A-mCherry-reverse-PGK-TetOn | This paper | N/A |
| pTRE-DRD2-P2A-mCherry-reverse-PGK-TetOn | This paper | N/A |
| pTRE-ARDB2-P2A-mCherry-reverse-PGK-TetOn | This paper | N/A |
| pTRE-GPR52-P2A-mCherry-reverse-PGK-TetOn | This paper | N/A |
| pTRE-HTR6-P2A-mCherry-reverse-PGK-TetOn | This paper | N/A |
| pTRE3G-DRD1-cHS4-EF1a-TetOn3G-P2A-mCherry-NLSx3 | This paper | N/A |
| pTRE3G-MC3R-cHS4-EF1a-TetOn3G-P2A-iRFP670 | This paper | N/A |
| pCMV- mouse PKAcat-linker-mCherry-CAAX | This paper | N/A |
| pCMV-mouse PKA-R1a-mEGFP | This paper | N/A |
| pCMV-mCerulean | This paper | N/A |
| pUCmini-iCAP-PHP.eB | Chan et al., 2017 | Plasmid #103005 |
| Software and algorithms |  |  |
| ImageJ (Fiji 1.48) | NIH | http://Fiji.sc |
| Python 3.8 |  | https://www.python.org |
| Other |  |  |
| Multiplate reader | TECAN | Spark |
| Confocal microscope | Carl Zeiss | LSM880 |
| Two-photon microscope | Olympus | FVMPE-RS |
| FASHIO-2PM | Ota et al., 2021 | N/A |
| Fiber photometry system | Doric | FPS_1S_GCaMP + Red Fluo |
| 400-μm-diameter mono fiberoptic cannula | Kyocera | N/A |

### RESOURCE AVAILABILITY

#### Lead contact

Further information and requests for resources and reagents should be directed to and will be fulfilled by the lead contact, Masayuki Sakamoto.

#### Materials availability

The cAMPinG1 and RCaMP3 sequence is available from GenBank (accession number: xxxxxxxx (cAMPinG1), and xxxxxxxx (RCaMP3)). Plasmids generated in this study have been deposited to RIKEN BRC (catalog number: RDBxxxxx-xxxxx). These will be available after the publication.

| Catalog number | Plasmid name |
| --- | --- |
| RDBxxxxx | pN1-cAMPinG1 |
| RDBxxxxx | pAAV-eSyn-cAMPinG1-NE-WPRE |
| RDBxxxxx | pAAV-eSyn-cAMPinG1-ST-WPRE |
| RDBxxxxx | pAAV-eSyn-RCaMP3-WPRE |

#### Data and code availability

- All data reported in this paper will be shared by the lead contact upon request.
- This paper does not report original code.
- Any additional information required to reanalyze the data reported in this paper is available from the lead contact upon request.

### EXPERIMENTAL MODEL AND SUBJECTIVE DETAILS

#### Animals

All animals were handled in accordance with the Kyoto University Guide, the University of Yamanashi Guide, the RIKEN guide, and the University of Tokyo Guide for the Care and Use of Laboratory Animals. Wild-type mice were group-housed and kept on a 12-h light/dark cycle with *ad libitum* food and water at room temperature. Wild-type animals used in this study were purchased from Japan SLC. Experiments were performed using male sex between 8 – 20 weeks of age.

#### Cell lines

HEK293T cells were obtained from the American Type Culture Collection (CRL-11268). Cells were cultured in Dulbecco’s Modified Eagle’s Medium (DMEM) (Nacalai) supplemented with 10% fetal bovine serum (FBS) (Sigma-Aldrich), 50 units/ml penicillin and 50 μg/ml streptomycin (Nacalai) at 37°C, 5% CO_2_ in a humidified atmosphere. *E. coli* DH5α and DH10B, and Stbl3 cells were obtained from Toyobo (DNA-9303), Invitrogen (18297010), and Invitrogen (C737303), respectively. Bacteria were incubated in Lysogeny Broth medium supplemented with antibiotics at 37°C.

### METHODS DETAILS

#### Plasmids

To develop cAMP sensors, PKA-R1α (amino acids 108-186 and 190-381) was obtained from a mouse cDNA library, and cpGFP and RSET domain were subcloned from GCaMP6f (Addgene plasmid # 52924). To enhance cytoplasmic localization of cAMPinG1, the F2A sequence of XCaMP-R was fused to the C-terminal of cAMPinG1 (Inoue et al., 2019). To develop soma-targeting cAMPinG1, the RPL10 domain was fused to the C-terminal of cAMPinG1 (Chen et al., 2020). For constructing RCaMP3, the cpRFP domain was synthesized (FragmentGENE, GENEWIZ), and RSET, M13, and CaM domains were obtained from jRGECO1a (Dana et al., 2016), F2A sequences were taken from XCaMP-R (Inoue et al., 2019). For site-directed mutagenesis, plasmid libraries were made using the inverse PCR method with PrimeSTAR Max DNA polymerase (Clontech), In-Fusion HD Cloning Kit (Clontech), and primers which included NNK codons, where K = G or T. For expression in *E. coli*, cAMP sensors were subcloned into a pBAD vector (Shen et al., 2018). For optogenetic stimulation, ChRmine-mScarlet-Kv2.1 was synthesized (FragmentGENE, GENEWIZ). For expression in HEK293T cells, Ca^2+^ or cAMP sensors were subcloned into a plasmid encoding CAG promoter and woodchuck hepatitis virus post-transcriptional regulatory element (WPRE). For expression of GPCRs-P2A-mCherry in HEK293T cells, pTRE3G-HA signal-Flag-GPCRs-P2A-mCherry-reverse-PGK-TetOn3G was made of Tet-ON 3G inducible expression system (Clontech) and PRESTO-Tango GPCR Kit (Kroeze et al., 2015).

The GenBank accession numbers for the sequence are xxxxxxxx (cAMPinG1) and xxxxxxxx (RCaMP3). cAMPinG1 and RCaMP3 plasmids were deposited to the RIKEN BRC (https://dna.brc.riken.jp/en/) for distribution following publication.

#### *In vitro* fluorometry for cAMP sensor screening

The plasmids for bacterial expression of cAMP sensors were transformed into *E. coli* strain DH10B (Invitrogen). *E. coli* cells were plated and cultured at 37 °C on Lysogeny Broth (LB) agar plate with ampicillin and 0.0004% arabinose. Each colony was used to inoculate 1.5 ml of LB liquid medium with ampicillin and 0.2% arabinose and grown at 37 °C overnight. After centrifugation, cells were resuspended in 150 µl suspension buffer (20 mM MOPS (pH 7.2), 100 mM KCl, 1 mM DTT, cOmplete EDTA free (Sigma-Aldrich)), sonicated at 4 °C, and centrifuged. The supernatant was collected. For fluorometry, the supernatant was diluted 20-fold with the suspension buffer. The diluted supernatant was applied to 96-well plates. cAMP (Tokyo Chemical Industry) was added to a final concentration of 300 µM for cAMP-saturated conditions. Fluorometric measurements were performed on a Spark microplate reader (TECAN) at room temperature by measuring the fluorescence intensity at the excitation wavelength of 485 nm, an excitation bandwidth of 20 nm, an emission wavelength of 535 nm, and an emission bandwidth of 20 nm.

#### *In vitro* cAMP fluorometry for HEK cell lysate

HEK293T cells were incubated at 37 °C on 6-well plates with 2 ml DMEM and 10% FBS. 1 µg DNA was transfected using X-tremeGENE HP DNA Transfection Reagent (Roche). Two days after the transfection, the cells were harvested to the 150 µl suspension buffer described above, sonicated at 4 °C, and centrifuged. The supernatant was collected. For cAMP-saturated conditions, the supernatant was diluted 20-fold with suspension buffer, and cAMP was added to a final concentration of 300 µM. Fluorometric measurements were performed at the excitation wavelength of 490 nm, excitation bandwidth of 20 nm, emission wavelength of 540 nm, and emission bandwidth of 20 nm. Excitation and emission spectra were taken at the emission wavelength of 555 nm and at the excitation wavelength of 460 nm, respectively. For measurement of cAMP affinity, the supernatant was diluted 40-fold with suspension buffer to a final concentration of 0, 3, 10, 30, 100, 300, 1,000, 3,000, 10,000, 30,000, 100,000, and 300,000 nM cAMP. For measurement of cGMP affinity, the supernatant was diluted 40-fold with suspension buffer to a final concentration of 0, 100, 300, 1,000, 3,000, 10,000, 30,000, 100,000, 300,000, and 1,000,000 nM cGMP (Abcam). The K_d_ value and Hill coefficient were calculated by fitting according to the Hill equation.

#### *In vitro* Ca^2+^ fluorometry for HEK cell lysate

Cell incubation, transfection, and collection was performed as described above. For Ca^2+^-saturated or Ca^2+^-free conditions, the supernatant was diluted 20-fold with the Ca^2+^-EGTA buffer (30 mM MOPS (pH 7.2), 100 mM KCl, 10 mM EGTA, 10 mM CaCl_2_, 1 mM DTT) or EGTA buffer (30 mM MOPS (pH 7.2), 100 mM KCl, 10 mM EGTA, 1 mM DTT), respectively. Fluorometric measurements were performed at the excitation wavelength of 560 nm, excitation bandwidth of 20 nm, emission wavelength of 610 nm, and emission bandwidth of 20 nm. Excitation and emission spectra were taken at the emission wavelength of 635 nm and at the excitation wavelength of 520 nm, respectively. For measurement of Ca^2+^ affinity, the supernatant was diluted 40-fold with a series of solutions with free Ca^2+^ concentration ranges from 0 nM to 3,900 nM (Zhao et al., 2011).

#### cAMP imaging in HEK293T cells

HEK293T cells were incubated at 37 °C in 35 mm glass bottom dishes or 96-well glass bottom plates. 1 µg DNA encoding the sensors was transfected as described above. One day after the transfection, the culture medium was replaced with Tyrode solution (129 mM NaCl, 5 mM KCl, 30 mM glucose, 25 mM HEPES-NaOH, pH 7.4, 2 mM CaCl_2,_ 2mM MgCl_2_). Imaging for cAMPinG1 was performed using LSM880 confocal microscope (Carl Zeiss). For time-lapse imaging, 405 nm and 488 nm wavelength lasers were used for excitation in turns. Forskolin (Nacalai) was added to a final concentration of 50 µM. For tetracycline-dependent expression of GPCRs-P2A-mCherry, doxycycline was added to a final concentration of 100 ng/ml 3 hours before the imaging. For single time-point imaging, the culture medium was replaced with Tyrode solution with or without drugs 20 minutes before the imaging. 405 nm and 488 nm wavelength lasers were used for ratiometric cAMP imaging, and 561 and 633 nm wavelength lasers was used for the visualization of GPCRs-expressing cells.

#### Two-photon red Ca^2+^ imaging in HEK293T cells

HEK293T cells were incubated at 37 °C in 35 mm glass bottom dishes. 0.8 µg DNA encoding the red Ca^2+^ sensors and pCMV-mCerulean was transfected as described above. One day after the transfection, the culture medium was replaced with Tyrode solution described above. Thirty seconds after bath application of ionomycin to a final concentration of 5 µM, two-photon imaging was performed with FVMPE-RS (Olympus) equipped with a water-immersion 25x objective (N.A.: 1.05, Olympus), a femtosecond laser (Insight DS+, Spectra-Physics), and two GaAsP detectors (Hamamatsu Photonics) with 495-540 nm and 575-645 nm emission filters (Olympus). Images (339 × 339 µm^2^, 1,024 × 1,024 pixels, single optical section) were collected. The laser was tuned to 880 nm for mCerulean and 1,040 nm at the front aperture of the objective) for the red Ca^2+^ sensors.

#### Stable cell lines generation

Stable cell lines were generated using lentiviral vectors. Lentiviral particles were produced by transfection of the packaging plasmids with polyethylenimine (PEI) into HEK293T cells using the same procedure as previously described (Imayoshi et al., 2013). Lentivirus-infected HEK293T cells were dissociated and isolated into multi-well plates. Single clones with bright fluorescence were picked, grown, and stored at -80 °C. To establish a triple stable cell line (**Figure 2**), cAMPinG1 single stable cell line was infected with lentivirus encoding TRE3G-DRD1-EF1a-TetOn3G-P2A-mCherry-NLSx3 and TRE3G-MC3R-EF1a-TetOn3G-P2A-iRFP670. (Shcherbakova et al., 2013)

#### Cell proliferation assay

The proliferation rate of the cAMPinG1 stable cell line was comparable with the original HEK293T cell line. The HEK293T cells were seeded on 6-well plates with 2 ml DMEM and 10% FBS or 1.5% FBS and incubated at 37 °C. The 1.5% FBS group was a positive control of slow cell proliferation. Twenty hours after the beginning of the culture, half of the wells were harvested, and the number of cells was counted using a counting chamber as a zero-time point. Sixty hours after the beginning of the culture, the other half of the wells were harvested and counted as a time point of 48 hours.

#### *In utero* electroporation

*In utero* electroporation (IUE) was performed as described previously with minor modifications (Sakamoto et al., 2022b). Briefly, ICR pregnant mice (Japan SLC) were anesthetized with an anesthetic mixture (0.075 mg/ml Medetomidine Hydrochloride, 0.40 mg/ml Midazolam, 0.50 mg/ml Butorphanol tartrate) and administered at 100 μl per 10 g of body weight (BW) intraperitoneally. Uterine horns were exposed. 2.0 µl of purified plasmid (1.0 µg/µl final concentration) was injected into the right lateral ventricle of embryos at embryonic day (E) 15. pCAG-cAMPinG1-NE and pCAG-RCaMP3-WPRE were delivered to induce the expression of cAMP and Ca^2+^ indicators in L2/3 pyramidal neurons in the V1. After soaking the uterine horn with warm saline (37 °C), each embryo’s head was carefully held between tweezers with platinum 5 mm disk electrodes (CUY650P5, Nepagene). Subsequently, five electrical pulses (45 V, 50 ms duration at 1 Hz) were delivered by an electroporator (NEPA21, Nepagene). After the electroporation, the uterine horns were returned into the abdominal cavity, and the skin was closed with sutures. Electroporated mice were used for cAMP and calcium imaging 4-10 weeks after birth and *in vivo* two-photon imaging.

#### AAV production and injection

Recombinant AAVs were produced using HEK293T cells as previously described with some modifications (Kawashima et al., 2013). The final titers were the followings: AAV2/1-CAG-DIO-cAMPinG1-NE (5.0 × 10^13^ GC/ml), AAV2/1-CAG-DIO-cAMPinG1mut-NE (2.0 × 10^13^ GC/ml), AAVPHP.eB-CaMKII-Cre (1.0 × 10^12^ GC/ml) for cAMP imaging in acute brain slice (**Figure S3**). AAV2/1-eSyn-jRGECO1a (3.0 × 10^13^ GC/ml), AAV2/1-eSyn-RCaMP3 (2.0 × 10^13^ GC/ml) for one-photon and two-photon calcium imaging in the barrel cortex (**Figures 3D-3J**). AAV2/1-eSyn-jRGECO1a (1.0 × 10^13^ GC/ml), AAV2/1-eSyn-RCaMP3 (1.0 × 10^13^ GC/ml) for two-photon mesoscale calcium imaging (**Figures 3K-3L**). AAV2/1-eSyn-cAMPinG1-ST (1.0 × 10^13^ GC/ml), AAV2/1-eSyn-RCaMP3 (1.0 × 10^13^ GC/ml) for two-photon imaging in the primary visual cortex (**Figures 4 and 5A-5G**). AAV2/1-hSyn-iCre (2.0 × 10^10^ GC/ml), AAV2/1-CAG-DIO-cAMPinG1-NE (3.0 × 10^13^ GC/ml), AAV2/1-CAG-DIO-RCaMP3 (1.0 × 10^13^ GC/ml, for infection marker), AAV2/1-CAG-DIO-ChRmine-mScarlet-Kv2.1 (1.0 × 10^12^ GC/ml) for cAMP imaging and optogenetic stimulation (**Figures 5H-5K**). AAV2/1-gfaABC1D-cAMPinG1-NE (1.0 × 10^13^ GC/ml), AAV2/1-gfaABC1D-RCaMP3 (1.0 × 10^13^ GC/ml) for astrocyte imaging (**Figure S6**). AAV2/1-eSyn-cAMPinG1-NE (7.0 × 10^12^ genome copies (GC)/ ml), AAV2/1-eSyn-RCaMP3 (1.0 × 10^13^ GC/ml) for fiber photometry in the dorsal striatum (**Figure 6**).

Stereotaxic virus injection was performed to C57BL/6N male mice aged 4-6 weeks anesthetized by the anesthetic mixture described above except for two-photon mesoscale imaging. A micropipette was inserted into the right primary visual cortex (A/P -3.85 mm, M/L +2.7 mm from the bregma, D/V -0.30 mm from the pial surface) or the barrel cortex (A/P -1.0 mm, M/L -3.0 mm from the bregma, D/V -0.20 mm from the pial surface). Then the virus solution of 500 - 800 nl was injected. Carprofen (5 mg/kg-BW; Zoetis) was administered intraperitoneally just after the injection experiment. Mice were subjected to imaging after 4-12 weeks of the injection.

#### cAMP imaging in acute brain slice

AAV (AAVPHP.eB-pCaMKII-Cre and AAV2/1-CAG-DIO-cAMPinG1, or AAV2/1-CAG-DIO-cAMPinG1mut) was injected into the primary visual cortex at a total volume of 500 nl (Chan et al., 2017). After 2 weeks of expression, mice were sacrificed by rapid decapitation after anesthesia with isoflurane. The brains were immediately extracted and immersed in gassed (95% O_2_/5% CO_2_) and ice-cold solution containing (in mM); (220 sucrose, 3 KCl, 8 MgCl_2_, 1.25 NaH_2_PO4, 26 NaHCO_3_, and 25 glucose). Acute coronal brain slices (280 μm thick) of the visual cortex were cut in gassed, ice-cold solution with a vibratome (VT1200, Leica, Germany). Brain slices were then transferred to an incubation chamber containing gassed artificial cerebrospinal fluid (ACSF) containing (in mM); 125 NaCl, 2.5 KCl, 1.25 NaH_2_PO_4_, 26 NaHCO_3_, 1 CaCl_2_, 2 MgCl_2_, 20 glucose at 34 °C for 30 minutes and subsequently maintained at room temperature before transferring them to the recording chamber and perfused with the ACSF solution described above, except using 2 mM CaCl_2_ and 1 mM MgCl_2_ at 30-32 °C. cAMP imaging was performed with an upright microscope (BX61WI, Olympus) equipped with an FV1000 laser-scanning system (FV1000, Olympus, Japan) and a 60 × objective lens (water-immersion, numerical aperture of 1.0, Olympus), a femtosecond laser (MaiTai, Spectra-Physics), and a GaAsP detector (Hamamatsu Photonics) with a 500-550 nm emission filter (Semrock). The laser wavelength was tuned at 940 nm (2 mW at the front aperture of the objective). Images (105.6 × 105.6 µm^2^, 640 × 640 pixels) were taken every 30 seconds. During the imaging, forskolin (Wako) and IBMX (Tocris) were added to a final concentration of 25 µM and 50 µM, respectively.

#### Simultaneous Ca^2+^ imaging and whole-cell recordings in acute brain slices

AAV (AAV2/1-eSyn-RCaMP3 or AAV2/1-eSyn-jRGECO1a) was injected into the barrel cortex (A/P -1.0 mm, M/L -3.0 mm from the bregma, D/V -0.2 mm from the pial surface) at 20 nl/min at a volume of 500 nl. After 4 weeks of expression, mice were sacrificed by rapid decapitation after anesthesia with pentobarbital (100 mg/kg). The brains were immediately extracted and immersed in gassed (95% O_2_/5% CO_2_) and ice-cold artificial cerebrospinal fluid (ACSF) containing (in mM); 124 NaCl, 2.5 KCl, 1.25 NaH_2_PO_4_, 26 NaHCO_3_, 2 CaCl_2_, 2 MgCl_2_, 10.1 glucose. Acute coronal brain slices (300 μm thick) of the barrel cortex were cut in gassed, ice-cold ACSF with a vibratome (VT1200S, Leica). Brain slices were then transferred to an incubation chamber containing gassed ACSF at 30°C for 60 minutes and subsequently maintained at room temperature before transferring them to the recording chamber at 35°C.

Whole-cell recordings were performed in the layer 2/3 pyramidal neurons of the barrel cortex with glass recording electrodes (5-8 MΩ) filled with the intracellular solution containing (in mM): 130 K-gluconate, 4 NaCl, 10 HEPES, 4 Mg-ATP, 0.3 Na-GTP, 7 dipotassium-phosphocreatine, pH adjusted to 7.0 with KOH (296 mOsm). Electrophysiological data were acquired using a patch-clamp amplifier (MultiClamp 700B, Molecular devices) filtered at 10 kHz and sampled at 20 kHz. Single action potentials were evoked by injecting a series of current pulses (2 ms duration) through the patch pipette. Each trial was repeated, and the mean value was presented.

Calcium imaging was performed using an upright microscope (BX51WI, Olympus) with a water immersion 40× (N.A.: 0.8) objective lens (Olympus). To acquire RCaMP3 and jRGECO1a images with LED light (MCWHLP1, Thorlabs), a U-MWIG3 fluorescence mirror unit (Olympus) was used. Fluorescent images were captured by a sCMOS camera (Orca-Flash 4.0 v3, Hamamatsu Photonics) controlled by HC Image software (Hamamatsu Photonics). Images were acquired at 50 Hz with 1 × 1 binning.

#### Simultaneous calcium imaging and cell-attached recordings *in vivo*

Loose-seal cell-attached recordings in vivo were performed as performed previously (Inoue et al., 2019). AAV (AAV2/1-eSyn-RCaMP3) was injected into the barrel cortex (A/P -1.0 mm, M/L - 3.0 mm from the bregma, D/V -0.2 mm from the pial surface) at 20 nl/min at a volume of 500 nl. After 4 weeks of expression, mice were head-fixed and anesthetized with isoflurane (1.5 ∼ 2.0 %) throughout the experiment, and body temperature was kept at 37°C with a heating pad. A craniotomy was made in the barrel cortex. The exposed brain was covered with 1.5% agarose in ACSF containing the following (in mM): 150 NaCl, 2.5 KCl, 10 HEPES, 2 CaCl_2_, 1 MgCl_2_, pH 7.3. A glass coverslip was then placed over the agarose to suppress the brain motion artifacts. A glass electrode (5-8 MΩ) was filled with ACSF containing Alexa 488 (200 µM). RCaMP3-expressing neurons were targeted using two-photon microscopy (Movable Objective Microscope, Sutter) with a tunable laser (InSight X3, Spectra-Physics) and a water-immersion 16 × (N.A.: 0.80) objective (Nikon). Fluorescence signals were collected using a GaAsP photomultiplier tube (Hamamatsu Photonics) with a 590-660 nm emission filter. After establishing the cell-attached configuration (20-100 MΩ seal), simultaneous spike recording and calcium imaging were performed at the soma (sampling rate = 30 Hz, 512 × 512 pixels). Electrophysiological data were acquired using a patch-clamp amplifier (MultiClamp 700B; Molecular devices) in current-clamp mode, filtered at 10 kHz, and sampled at 20 kHz. The laser was tuned to 1,040 nm (40 mW at the front aperture of the objective).

#### Fiber photometry

Mice were anesthetized by the anesthetic mixture described above. Then, a 400-µm-diameter mono fiberoptic cannula (Kyocera) was implanted above the right dorsal striatum. A custom-made metal headplate was attached to the skull with dental cement. Mice were subjected to imaging after more than 2 days of the surgery.

Dual-color fiber photometry for cAMPinG1 and RCaMP3 was performed using GCaMP & Red Fluorophore Fiber Photometry System (Doric) with 405 nm, 470 nm, and 560 nm LED and 400-µm-diameter 0.57-N.A. Mono Fiber-optic Patch Cords (Doric). cAMPinG1 was excited by 405 nm and 470 nm LED, and its fluorescence was detected with a single photodetector with a 500-540 nm emission filter. The green fluorescence by 470 nm excitation was demodulated from green fluorescence by 405 nm excitation by lock-in amplifier detection. Simultaneously, RCaMP3 was excited by 560 nm LED and spectrally distinguished from cAMPinG1 signals by a dichroic mirror, and its fluorescence was detected with a single photodetector with a 580-680 nm emission filter. Photometry data was recorded at a sampling rate of 30 Hz. Mice were head-fixed during the recordings.

#### Cranial window implantation

Craniotomy was performed as described previously (Sakamoto et al., 2022a). Mice were anesthetized by the anesthetic mixture described above. Before surgery, dexamethasone sodium phosphate (2 mg/kg-BW; Wako) and carprofen (5 mg/kg-BW; Zoetis) were administered to prevent inflammation and pain. During surgery, mice were put on a heating pad, and body temperature was kept at 37 °C. A custom-made stainless head plate was fixed to the skull using cyanoacrylate adhesive and dental cement (Sun-medical) above the right visual cortex. A craniotomy was drilled with a 2.5 mm diameter, and the brain was kept moist with saline. A cover glass (3 mm diameter, #0 thickness, Warner Instruments) was placed over the craniotomy site with surgical adhesive glue (Aron Alpha A, Sankyo). The mice were subjected to imaging more than 18 h after the surgery.

#### *In vivo* two-photon imaging

*In vivo* two-photon imaging was performed with FVMPE-RS (Olympus) equipped with a water-immersion 25x objective (N.A.: 1.05, Olympus), a femtosecond laser (Insight DS+, Spectra-Physics), and two GaAsP detectors (Hamamatsu Photonics) with 495-540 nm and 575-645 nm emission filters (Olympus). For somatic cAMP imaging of cAMPinG1 and cAMPinG1mut expressed by *in utero* electroporation, images (339 × 339 µm^2^, 512 × 512 pixels, single optical section) were collected at 15 Hz in the awake condition. The laser was tuned to 940 nm (48.6 mW at the front aperture of the objective). For cAMPinG1-NE spine imaging, images (28.8 × 38.4 µm^2^, 96 × 128 pixels, single optical section) were collected at 7.5 Hz in the condition anesthetized lightly by isoflurane (0.5% v/v). The laser power was 23.5 mW at the front aperture of the objective. For RCaMP3 and cAMPinG1-ST imaging, sequential excitation at 940 nm and 1,040 nm was used for dual-color imaging. Images (339 × 339 µm^2^, 512 × 512 pixels) with three optical planes with plane spaced 30 µm apart in depth) were collected at 3.4 Hz per plane using a piezo objective scanner (Olympus). The laser power for 940 nm excitation was set to 47.6 mW, and for 1,040 nm excitation it was set to 113.7 mW. The imaging with visual stimuli (**Figures 5A-5G**) were followed by the imaging during forced running (**Figure 4**) on the same cell population. For cAMP and Ca^2+^ imaging in astrocytes, images (288 × 384 µm^2^, 192 × 256 pixels) were collected at 1.6 Hz in the awake condition. The laser power for 940 nm excitation was set to 26.5 mW, and for 1,040 nm excitation, it was set to 52.2 mW. For single-cell cAMP imaging with optical stimulation using soma-targeted ChRmine, images (86.4 × 115.2 µm^2^, 192 × 256 pixels) were collected at 3.2 Hz in the awake condition with 940 nm excitation. 1,040 nm two-photon excitation was employed for 4-second spiral scanning with a diameter of 11 µm. The imaging with a 940 nm laser was temporally stopped during the optical excitation. The laser power for 940 nm excitation was set to 4.1 mW, and for 1,040 nm photostimulation, it was set to 39.8 mW.

#### Large field of two-photon mesoscale imaging

AAV (AAV2/1-eSyn-RCaMP3) was injected into the neonatal somatosensory cortex as previously described (Oomoto et al., 2021). After 8 weeks of AAV injection, a 4.5-mm diameter craniotomy was performed over an area including the primary somatosensory area of the right hemisphere. A head plate was also fixed to the skull above the cerebellum.

Two-photon imaging was performed with FASHIO-2PM (Ota et al., 2021) equipped with a femtosecond laser (Chameleon Discovery, Coherent). The laser wavelength was 1,040 nm. The field-of-view was 3.0 × 3.0 mm^2^ (2,048 × 2,048 pixels). The sampling rate was 7.5 Hz. Laser power of 270 and 360 mW at the front of the objective lens was used to observe layer 2/3 and 5 neurons of awake mice, respectively.

#### Physical stimulation

Airpuff stimuli (2Hz, 0.1 s duration, 40 times) were generated using a microinjector (BEX). For the forced running task, mice were head-fixed, and a custom-made treadmill was turned on during recordings. Moving grating stimuli were generated using the Psychopy in Python as described previously with some modifications (Chen et al., 2013). The gratings were presented with an LCD monitor (19.5 inches, 1,600 × 900 pixels, Dell), placed 25 cm in front of the center of the left eye of the mouse. Each stimulus trial consisted of a 4-s blank period (uniform grey at mean luminance) followed by a 4-s drifting sinusoidal grating (0.04 cycles per degree, 2 Hz temporal frequency). Eight drifting directions (separated by 45°, in order from 0° to 315°) were used. The timing of each moving grating stimulus and the initiation of imaging were monitored with a data acquisition module (USB-6343, National Instruments).

### QUANTIFICATION AND STATISTICAL ANALYSIS

#### *In vitro* fluorometry

*In vitro* fluorometry analysis was performed using Python (https://www.python.org) and Excel (Microsoft). For both Ca^2+^ and cAMP sensors, the K_d_ value and Hill coefficient were calculated by fitting according to the Hill equation.

#### Image analysis

Image analyses were performed with ImageJ (NIH) and Python. For somatic cAMP imaging of cAMPinG1 and cAMPinG1mut expressed by *in utero* electroporation (**Figures 2A-2E**) and simultaneous RCaMP3 and cAMPinG1-ST imaging (**Figures 4 and 5A-5G**), motion correction was performed with Suite2p toolbox (https://github.com/MouseLand/suite2p) (Pachitariu et al., 2016). For cAMPinG1-NE spine imaging (**Figures 2G-2I**), mesoscale RCaMP3 imaging (**Figures 3K-3L and S5**), Ca^2+^ and cAMP imaging in astrocytes (**Figure S6**), and cAMP imaging with optical stimulation using soma-targeted ChRmine (**Figures 5H-5K**), motion correction was performed with TurboReg (Thevenaz et al., 1998).

Regions of interest (ROI) detection for *in vivo* Ca^2+^ imaging was performed with Suite2p. ROI detection for HEK cell live imaging and *in vivo* cAMP imaging were performed with ImageJ (NIH) and Cellpose (Stringer et al., 2021). ROIs for cAMPinG1-NE spine imaging (**Figures 2G-2I**), Ca^2+^ and cAMP imaging in astrocytes (**Figure S6**), and cAMPinG1-NE imaging with soma-targeted ChRmine (**Figures 5H-5K**) were drawn manually. The dynamic range was calculated as ΔF/F = (F – F_0_)/F_0,_ where F_0_ was the average fluorescence intensity before stimulations after the subtraction of background fluorescence. No bleaching correction was performed in any analyses except Ca^2+^ imaging in acute brain slices (**Figure 3**). No fluorescence crosstalk correction was performed.

For cAMP imaging using acute brain slices (**Figure S3**), background subtraction was performed before calculating ΔF/F. ROIs were manually selected around somata in the time series averaged image. ΔF/F was calculated as (F-F_0_) / F_0_, where F is the fluorescence intensity at any time point and F_0_ is the average fluorescence before the drug application. For *in vivo* cAMPinG1-NE spine imaging (**Figures 2G-2I**), the period after stimulation was defined as a 15 s period starting 10 s after the end of airpuff stimulation. For two-photon Ca^2+^ imaging in HEK293T cells (**Figure S4**), ROIs drawn based on mCerulean images with Cellpose were used for both Ca^2+^ sensor and mCerulean images. The red fluorescence intensity was divided by mCerulean fluorescence intensity in each cell for normalization. For Ca^2+^ imaging using acute brain slices (**Figures 3D-3H**), background subtraction and bleach correction were performed before calculating ΔF/F. ROIs were manually selected around somata in the time series averaged image. ΔF/F was calculated as (F-F_0_) / F_0_, where F is the fluorescence intensity at any time point and F_0_ is resting baseline fluorescence measured 200 ms before stimulation. The peak amplitude was defined as the maximum value of ΔF/F after the stimuli. The rise and decay curves were fit to a single exponential. The rise time was defined as the time from the beginning of the stimulus to the time point of the peak fluorescence amplitude. The half-decay time was defined as the time from the maximum value of ΔF/F to half of that value. For simultaneous RCaMP3 and cAMPinG1-ST imaging (**Figures 4 and 5A-5G**), ROIs for RCaMP3 and cAMPinG1-ST were drawn independently using Suite2p and Cellpose, respectively. The cells that had ROIs for both RCaMP3 and cAMPinG1-ST were selected for further analysis. Because the imaging using visual stimulus (**Figures 5A-5G**) was followed by the imaging using forced running (**Figure 4**) on the same cell population, the same ROIs were used for both analyses. Motion-related neurons were defined as neurons that showed Ca^2+^ ΔF/F more than 0.3 during the running period (**Figure 4**). Ca^2+^ responses to 4 s visual stimulation were defined as averaged ΔF/F during the 4 s stimulation. cAMP responses to 4 s visual stimulation were defined as averaged ΔF/F during 2 s periods that started 2 s after the end of the visual stimulus (**Figures 5A-5D**). Neurons that responded to repetitive 8 s visual stimuli were defined as neurons that showed Ca^2+^ ΔF/F more than 0.3 during the stimulation period (**Figures 5E-5G**). The period after repetitive visual stimuli was defined as a 40 s period starting after the end of the stimuli. The direction selectivity index (DSI) was calculated for cells showing Ca^2+^ responses. The preferred direction (θ_pref_) of each cell was defined as the stimulus that induced the largest Ca^2+^ ΔF/F. The DSI was defined as DSI = (R_pref_ - R_pref+_π)/(R_pref_ + R_pref+_π), where R_pref_ and R_pref+_π are ΔF/F at the preferred (θ_pref_) and the opposite direction (θ_pref_ +π) respectively. Imaging frames with significant motion artifacts are removed and supplied with the preceding frames. For cAMPinG1-NE imaging with soma-targeted ChRmine (**Figures 5K**), cAMP ΔF/F was defined as averaged ΔF/F during a 10 s period starting 20 s after the end of optical stimulation.

#### Fiber photometry

Fiber photometry analysis was performed using Python. The cAMPinG1 signal was calculated as follows: (470 nm signal) / (405 nm signal). The 560 nm signal was recognized as the RCaMP3 signal. The period during stimulation was defined as a 10 s running period, and the period after stimulation was defined as 10 s after the end of the running period (**Figure 6D**).

#### Statistical analysis

All statistical analyses of the acquired data were performed with Python. For each figure, a statistical test matching the structure of the experiment and the structure of the data was employed. All tests were two-tailed. \**p* < 0.05; \*\**p* < 0.01; \*\*\**p* < 0.001; NS, not significant (*p* > 0.05) for all statistical analyses presented in figures. No statistical tests were done to predetermine the sample size. Data acquirement and analysis were not blind. Experimental sample sizes are mentioned in the figure panel and legends.

## ACKNOWLEDGMENTS

We thank the following researchers for kindly sharing their reagents; Haruhiko Bito (eSyn promoter, cHS4, XCaMP-R, and pCAG-Flpo plasmids), Naoto Saito (gCarvi plasmid), Tyler Jacks (Addgene plasmid #12093), Vladislav Verkhusha (Addgene plasmid #45457), Jinhyun Kim (Addgene plasmid # 51904), Baljit Khakh (Addgene plasmid #52924), Tetsuya Kitaguchi (Addgene plasmid #73938), Douglas Kim (Addgene plasmid #100854), Viviana Gradinaru (Addgene plasmid #103005), Robert Campbell (Addgene plasmid #105864), Jennifer Garrison (Addgene plasmid #158777), Bryan Roth (Addgene kit #1000000068). We also thank Yu Kato for his technical assistance and the Sakamoto and Imayoshi laboratory members for their support and comments.

This work was supported in part by grants from Precursory Research for Embryonic Science and Technology (PRESTO)-JST (JPMJPR1906 to M.S.), ACT-X-JST (JPMJAX211K to T.Y.), Grant-in-Aid for Brain Mapping by Integrated Neurotechnologies for Disease Studies (Brain/MINDS) (JP22dm0207090 to I.I., JP19dm0207069 to S.Y., JP21dm0207001 to M.M., JP20dm020706 to M.S.), Strategic Research Program for Brain Sciences (JP22wm0525018 to S.Y., JP22wm0525004 to M.S.), Interstellar Initiative Beyond (JP22jm0610068 to M.S.), JSPS KAKENHI (JP21K15207 to T.Y., JP20H05775 to M.M., JP21K19429, JP20H04122 to M.S.), RIKEN Special Postdoctoral Researchers Program (to H.U.), Takeda Science Foundation (to M.S.), Lotte Foundation (to M.S.), The Konica Minolta Science and Technology Foundation (to M.S.), Tokyo Biochemical Research Foundation (to M.S.), Brain Science Foundation (to M.S.).

## AUTHOR CONTRIBUTIONS

T.Y. designed and developed cAMP sensors. T.Y. and M.S. designed and developed the RCaMP3. T.Y. performed most of the experiments and analyzed data. S.M. and K.K. performed electrophysiological recordings for RCaMP3 characterization. U.H. and M.M. performed Ca^2+^ imaging with the FASHIO-2PM. M.T. and S.Y. performed pharmacological experiments for cAMPinG1 in acute brain slices. T.Y. and M.S. wrote the paper with input from all authors. I.I. and M.S. supervised the entire project.

