## Supplementary Information for "A multicolor suite for deciphering population coding in calcium and cAMP *in vivo*"

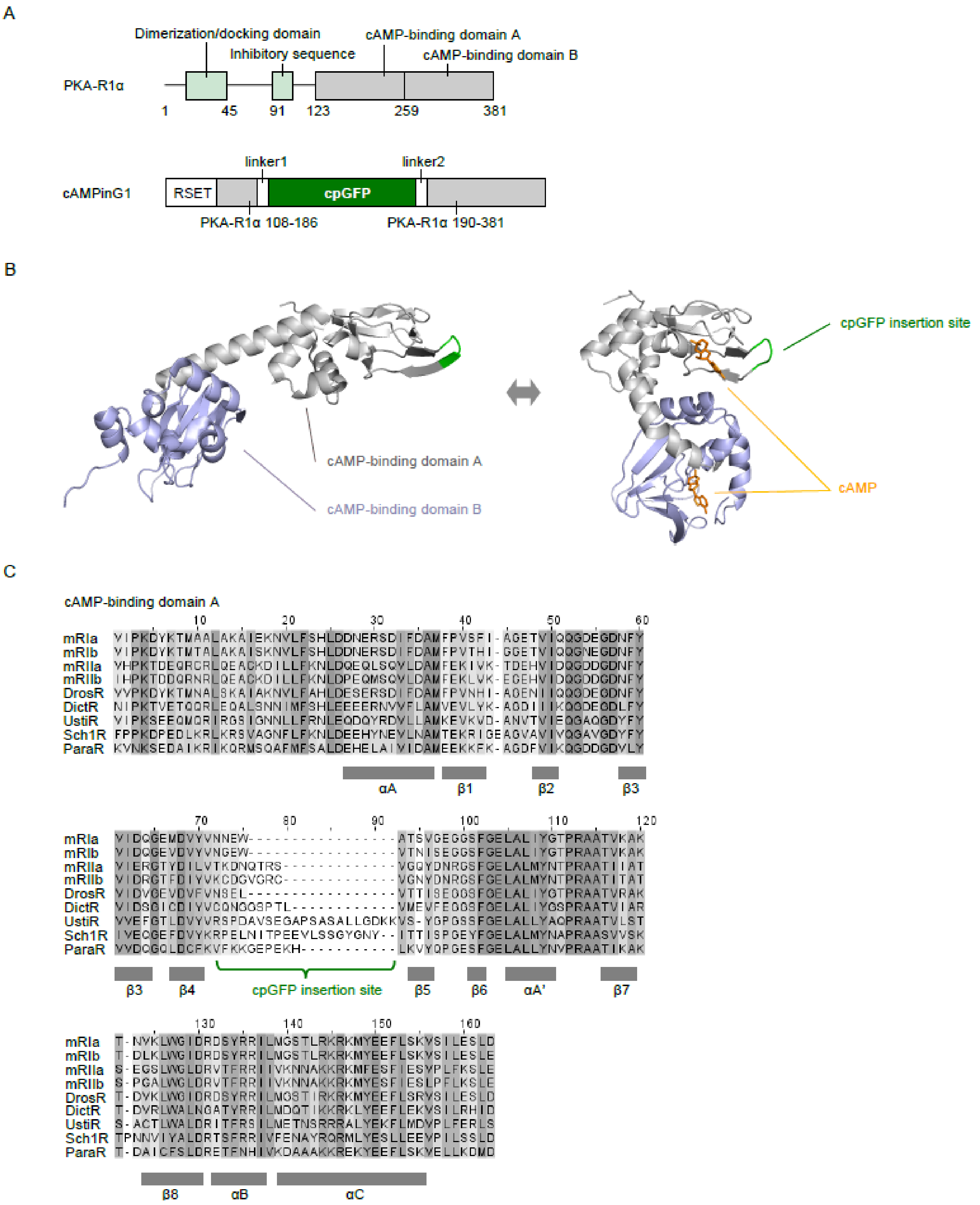


**Figure S1. Design of cAMPinG1, related to Figure 1**

**(A)** Primary structures of mouse PKA regulatory-subunit 1α (top) and cAMPinG1 (bottom). **(B)** Tertiary structures of cAMP-free (left, PDB: 2QCS, catalytic-subunit is hidden) and cAMP-binding (right, PDB: 1RGS) PKA R1α. **(C)** The amino acid sequence of cAMP-binding domains of PKA regulatory-subunit. The cpGFP insertion site of cAMPinG1 is indicated in a green bracket. mR1a, mouse PKA-RIa; mR1b, mouse PKA-R1b; mR2a, mouse PKA-R2a; mR2b, mouse PKA-R2b; DrosR, Drosophila melanogaster PKA-R; DictR, Dictyostelium discoideum PKA-R; UstiR, Ustilago maydis PKA-R; Sch1R, Schizosaccharomyces pombe PKA-R; ParaR, Paramecium tetraurelia PKA-R.

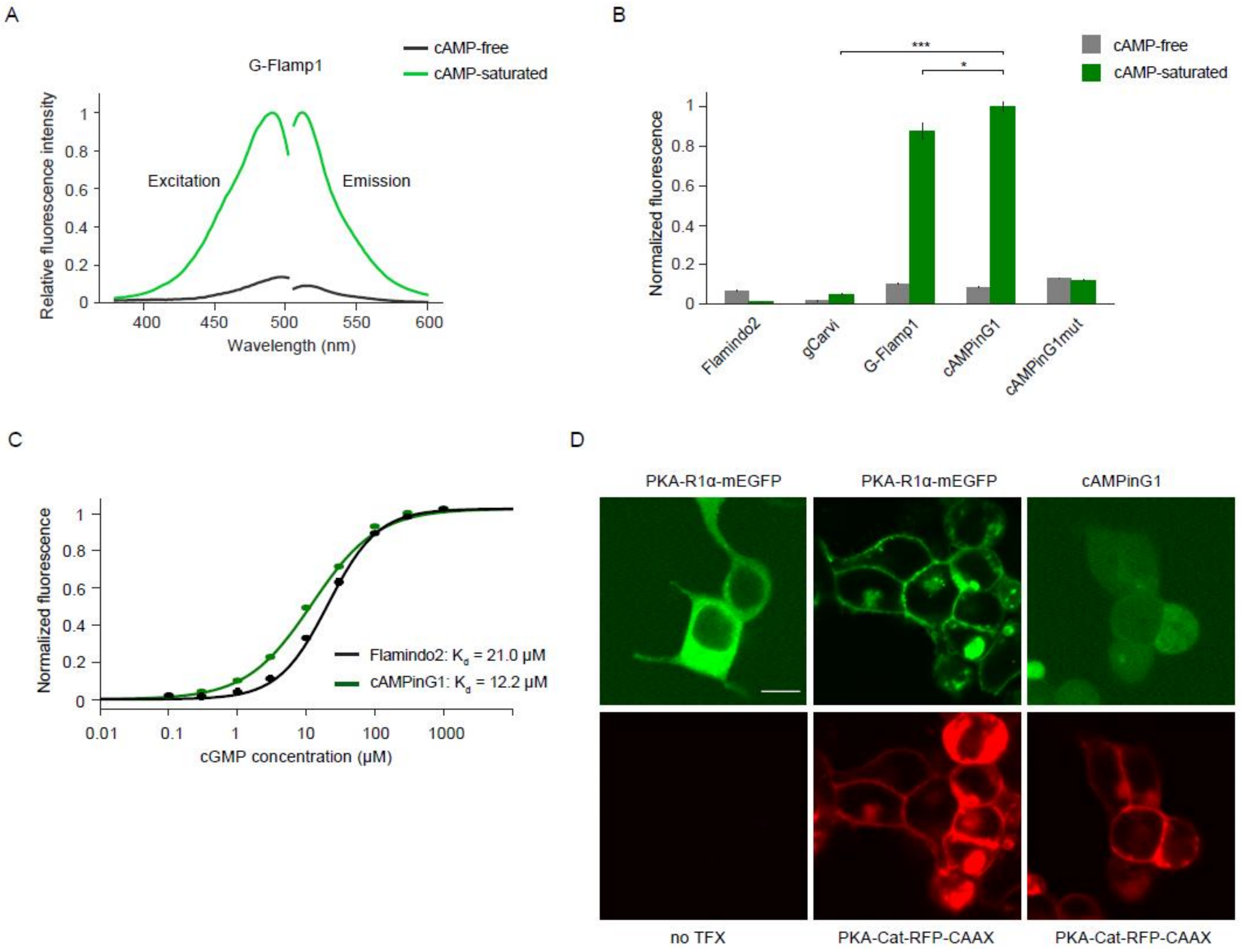


**Figure S2. Characterization of cAMP sensors *in vitro*, related to Figure 1**

**(A)** Excitation and emission spectra of G-Flamp1 in cAMP-free (black) and cAMP-saturated (light green) states. **(B)** Fluorescence intensities of cAMP sensors in cAMP-free and cAMP-saturated states in HEK293T cell lysate. n = 4 wells in each sensor. Tukey's post hoc test following one-way ANOVA. **(C)** cGMP titration curves of Flamindo2 and cAMPinG1. Flamindo2: cGMP $K_d$ = 21.0 μM, cAMPinG1: cGMP $K_d$ = 12.2 μM. n = 4 wells (Flamindo2), n = 4 wells (cAMPinG1). **(D)** Binding assay of PKA-R1α-mEGFP and cAMPinG1 with PKA-catalytic subunit-RFP-CAAX. Scale bar, 10 μm. All error bars denote the SEM.

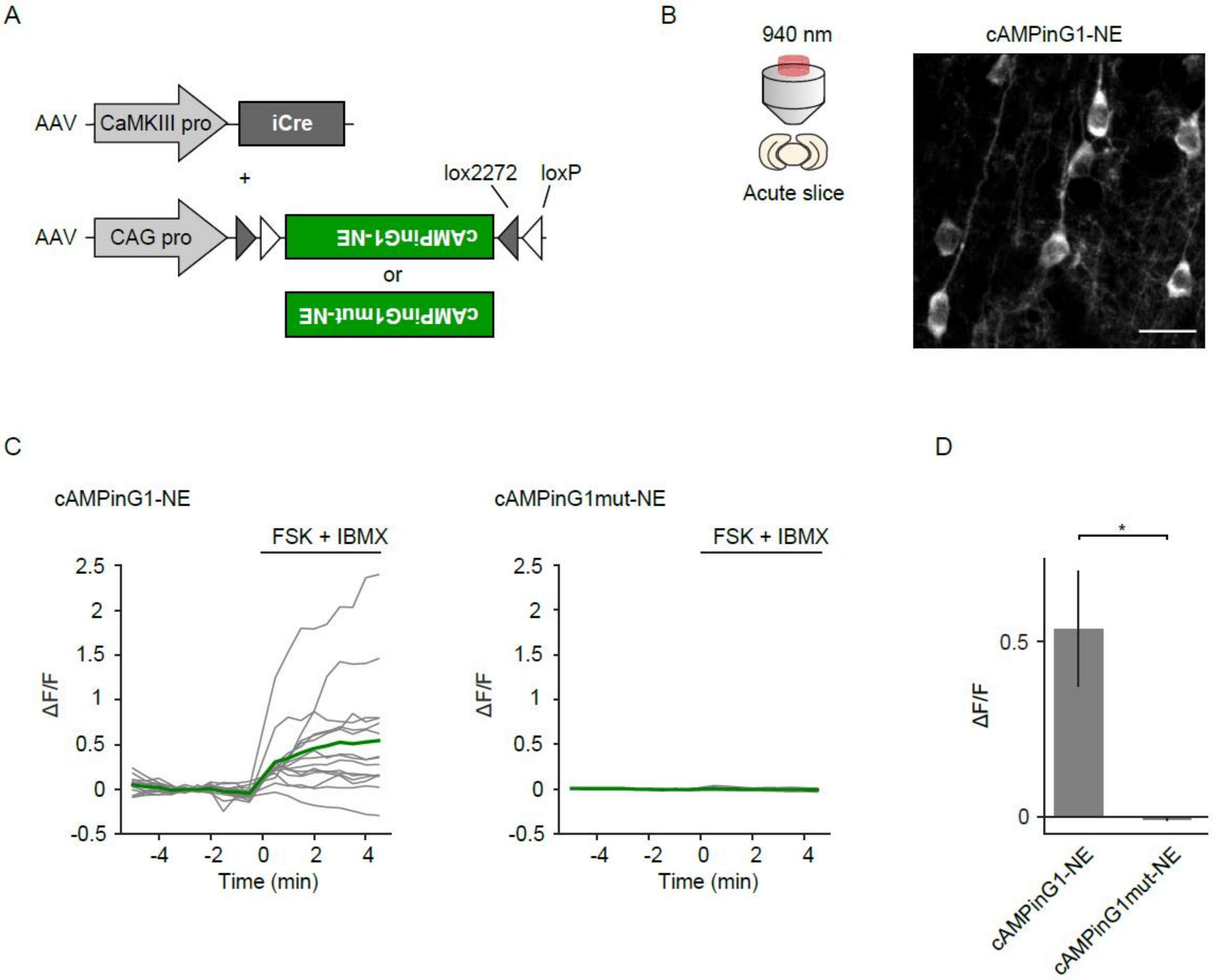


**Figure S3. Characterization of cAMPinG1 in acute brain slices, related to Figure 2**

**(A)** Schematic of AAVs for sparse expression of cAMPinG1-NE and cAMPinG1mut-NE. **(B)** Schematic of imaging settings (left). A representative two-photon image of cAMPinG1 in an acute brain slice (right). Scale bar, 20 μm. **(C)** Traces of cAMPinG1-NE (left) and cAMPinG1mut-NE (right) in response to application of forskolin and IBMX to a final concentration of 25 μM and 50 μM, respectively. Grey lines denote individual trace, and colored thick lines denote average response. The black vertical lines indicate stimuli. n = 16 neurons in 5 slices (cAMPinG1-NE), 12 neurons in 2 slices (cAMPinG1mut-NE). **(D)** Averaged ΔF/F of cAMPinG1-NE and cAMPinG1mut-NE in response to forskolin application. n = 16 neurons in 5 slices (cAMPinG1-NE), n = 12 neurons in 2 slices (cAMPinG1mut-NE). Unpaired t-test. Error bars denote the SEM.

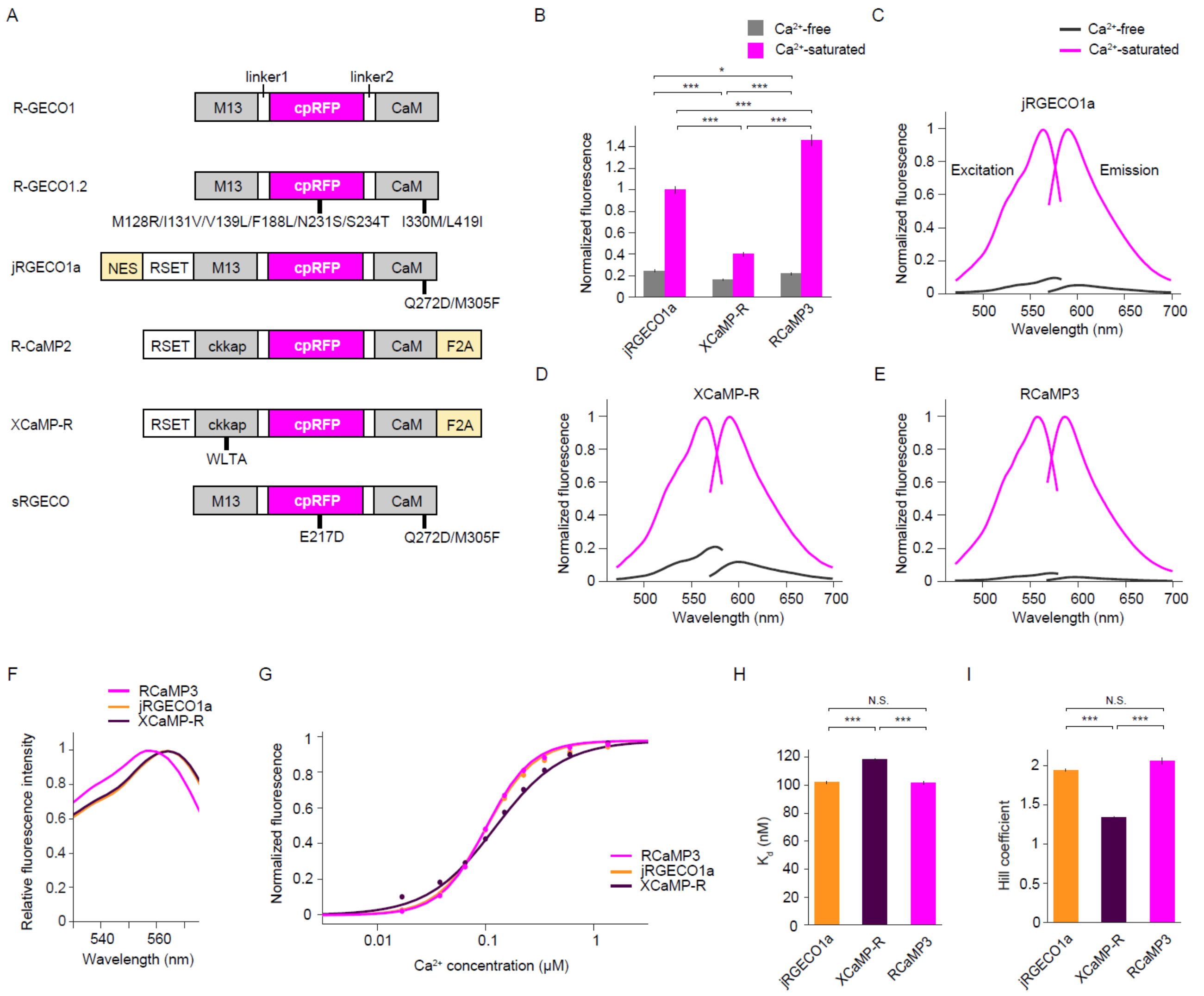


**Figure S4. Properties of red $Ca^{2+}$ sensors, related to Figure 3**

**(A)** Primary structures of red $Ca^{2+}$ sensors. Mutations are indicated in R-GECO1 numbering. **(B)** Fluorescence intensities of red $Ca^{2+}$ sensors in $Ca^{2+}$-free and $Ca^{2+}$-saturated conditions in HEK cell lysate. n = 4 wells (jRGECO1a), n = 4 wells (XCaMP-R), n= 4 wells (RCaMP3). Tukey's post hoc test following one-way ANOVA. **(C-E)** Excitation and emission spectra of jRGECO1a, XCaMP-R, and RCaMP3 in $Ca^{2+}$-free (dark magenta) and $Ca^{2+}$-saturated (light magenta) states. n = 4 wells (jRGECO1a), n = 4 wells (XCaMP-R), n= 4 wells (RCaMP3). **(F)** Comparison of excitation spectra of red $Ca^{2+}$ indicators. RCaMP3 was blue-shifted. **(G-I)** $Ca^{2+}$ titration curves (G), $K_d$ values (H), and hill coefficients (I) of red $Ca^{2+}$ sensors. n = 4 wells (jRGECO1a), n = 4 wells (XCaMP-R), n= 4 wells (RCaMP3). Tukey’s post hoc test following one-way ANOVA. All error bars denote the SEM.

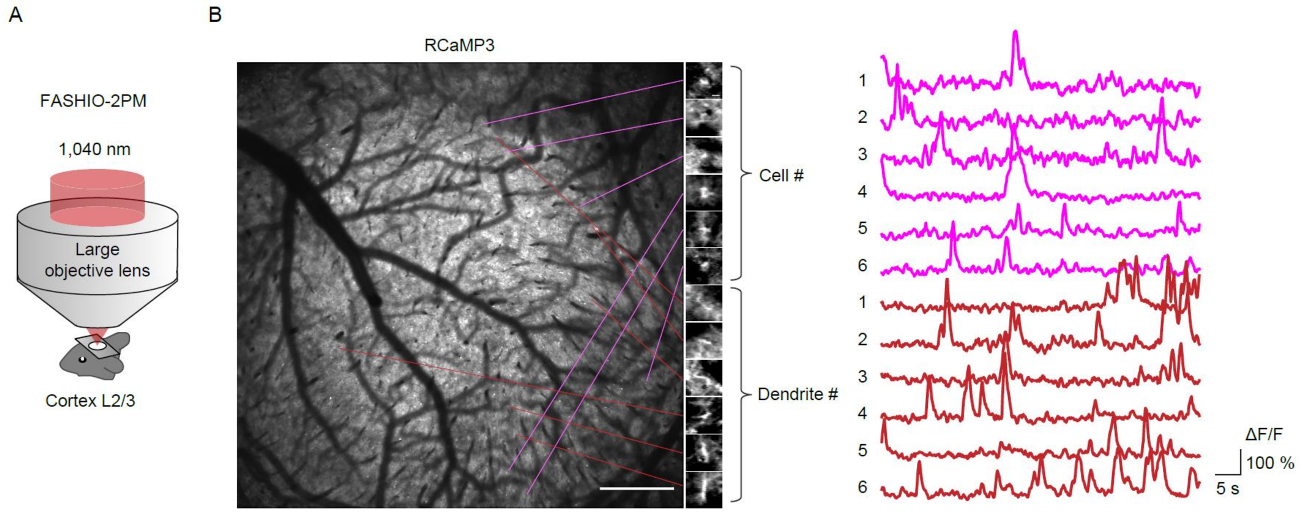


**Figure S5. Two-photon mesoscale RCaMP3 imaging in layer 2/3, related to Figure 3**

**(A)** Schematic of the experimental procedure of two-photon mesoscale $Ca^{2+}$ imaging using fast-scanning high optical invariant two-photon microscopy (FASHIO-2PM). Cortical layer 2/3 (L2/3) neurons in the field-of-view (FOV, 3.0 × 3.0 $mm^2$) were imaged by 1,040 nm excitation. **(B)** Left: A representative full FOV of FASHIO-2PM. Scale bar, 500 µm. Right: Magnified images and $Ca^{2+}$ traces of representative 6 somata and 6 dendrites. Scale bar, 10 µm.

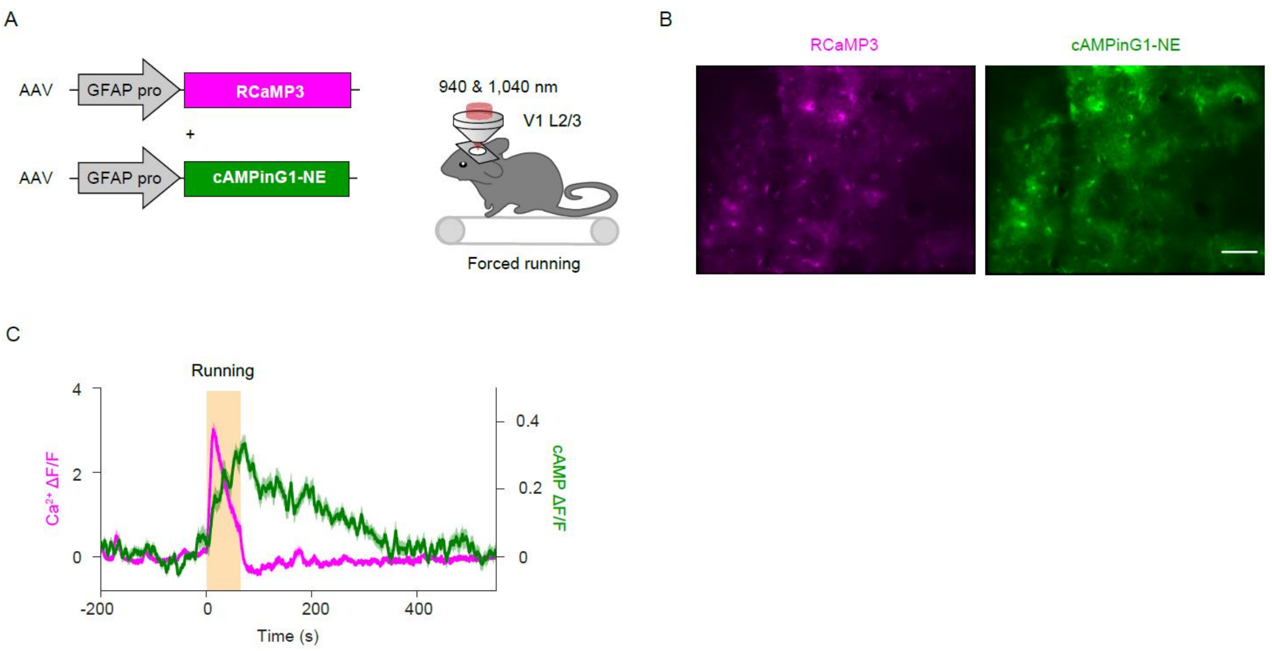


**Figure S6. $Ca^{2+}$ and cAMP imaging of astrocytes during forced running, related to Figure 4**

**(A)** Schematic of the experimental procedure. The GFAP promoter was used to express RCaMP3 and cAMPinG1-NE in astrocytes. **(B)** Representative images of cAMPinG1 and RCaMP3. Scale bar, 50 µm. **(C)** Averaged fluorescence transients of cAMPinG1 (green) and RCaMP3 (magenta). n = 63 cells in 3 mice. Shaded areas denote the SEM.

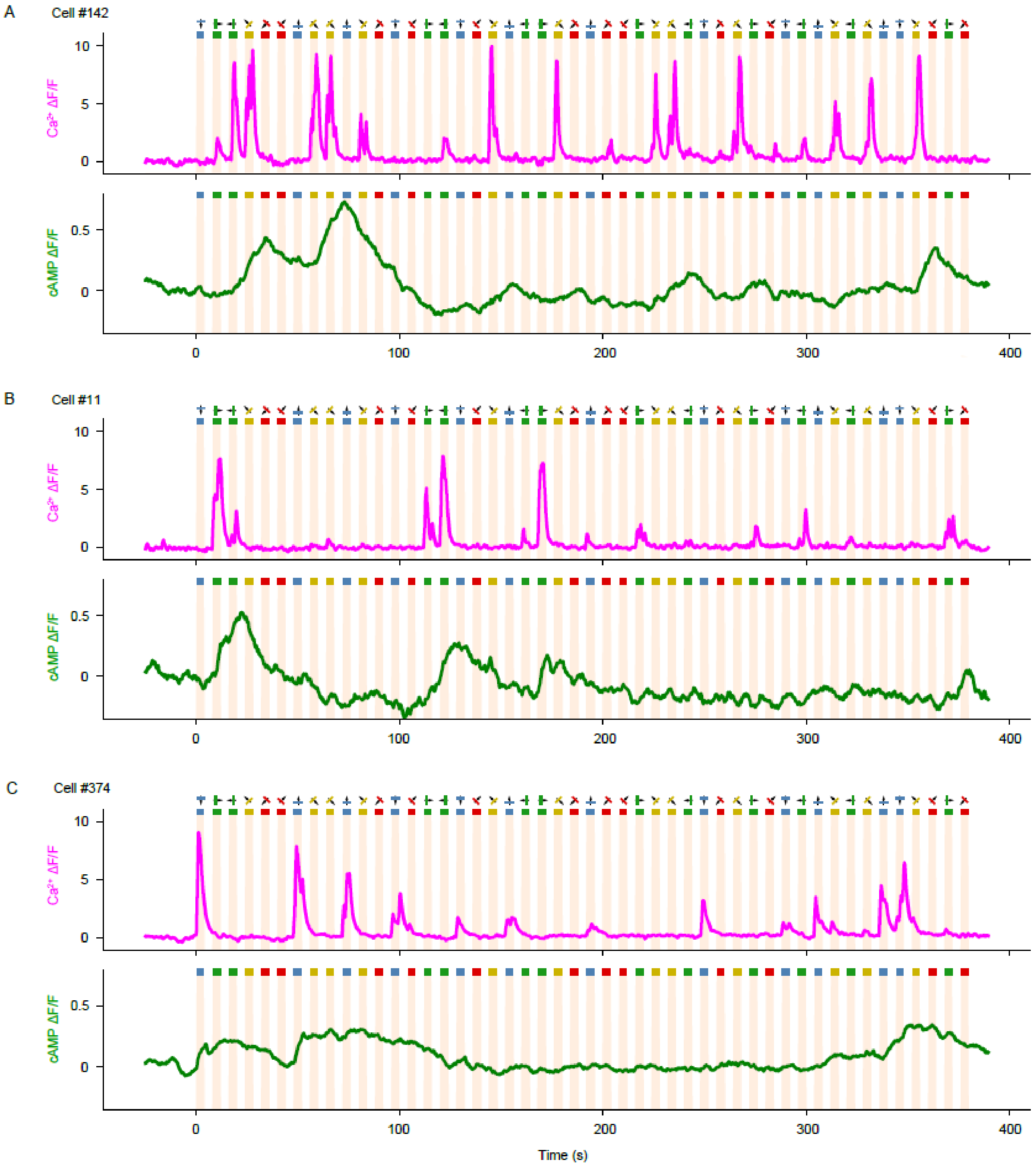


**Figure S7. Raw traces of $Ca^{2+}$ and cAMP in representative cells during visual stimuli, related to Figure 5**

**(A-C)** $Ca^{2+}$ and cAMP traces of representative cells in Figures 5B-6D.

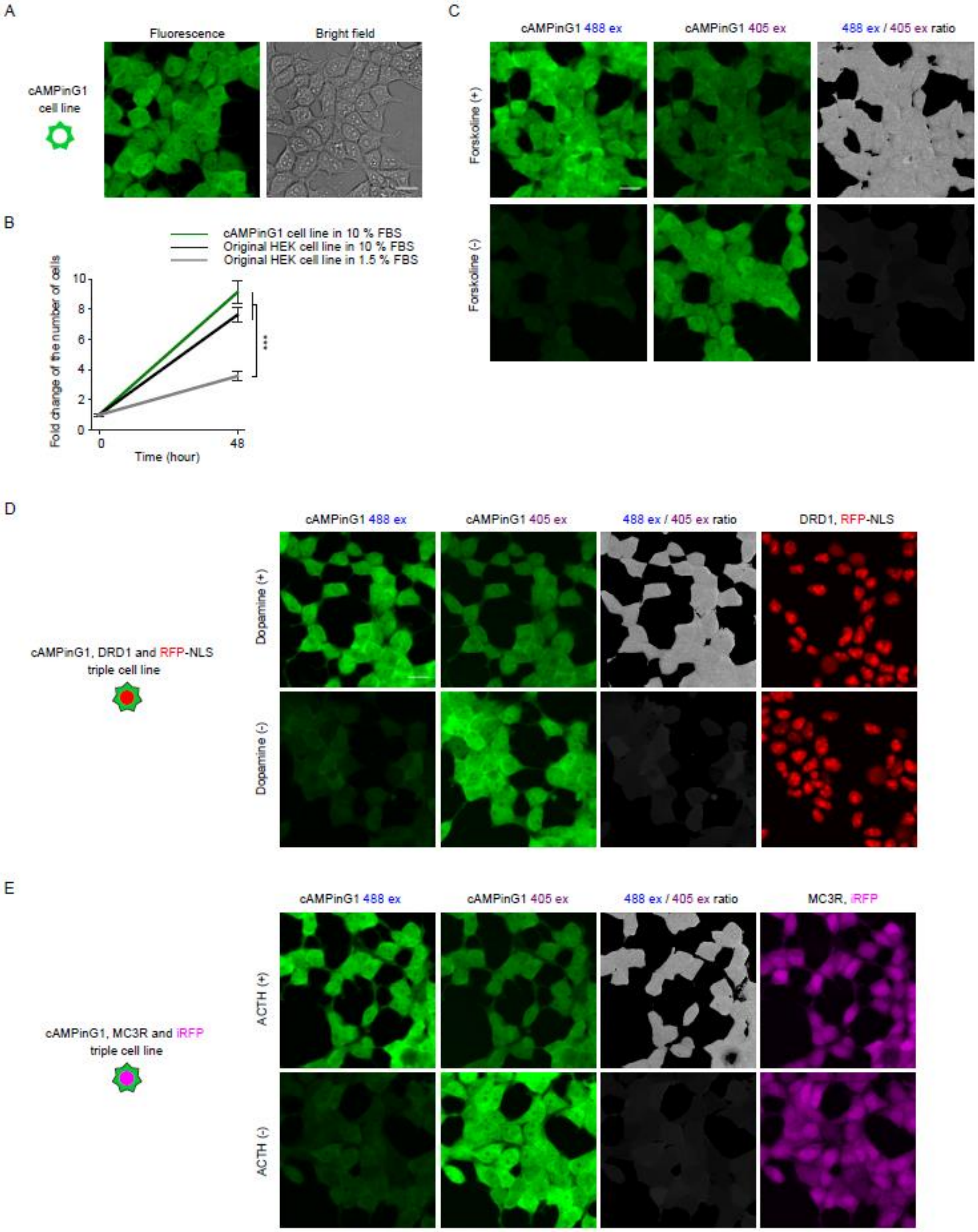


**Figure S8. Establishing a stable cell line of cAMPinG1, related to Figure 7**

**(A)** A representative image of the cAMPinG1 stable cell line. Scale bar, 20 μm. **(B)** Proliferation assay of the cAMPinG1 stable cell line and original HEK cell line. The original HEK cell line in 1.5 % FBS condition was for a negative control of the assay. n = 6 wells in each cell line. Tukey's post hoc test following one-way ANOVA. Error bars denote the SEM. **(C)** Single timepoint imaging of cAMPinG1 cell line in the absence (top) or presence (bottom) of 50 μM forskolin. Scale bar, 20 μm. **(D)** Single timepoint imaging of the cAMPinG1, DRD1, and mCherry triple stable cell line in the absence (top) and presence (bottom) of 100 nM dopamine. Scale bar, 20 μm. **(E)** Single timepoint imaging of the cAMPinG1, MC3R and iRFP670 triple stable cell line in the absence (top) or presence (bottom) of 1,000 nM ACTH. Scale bar, 20 μm.

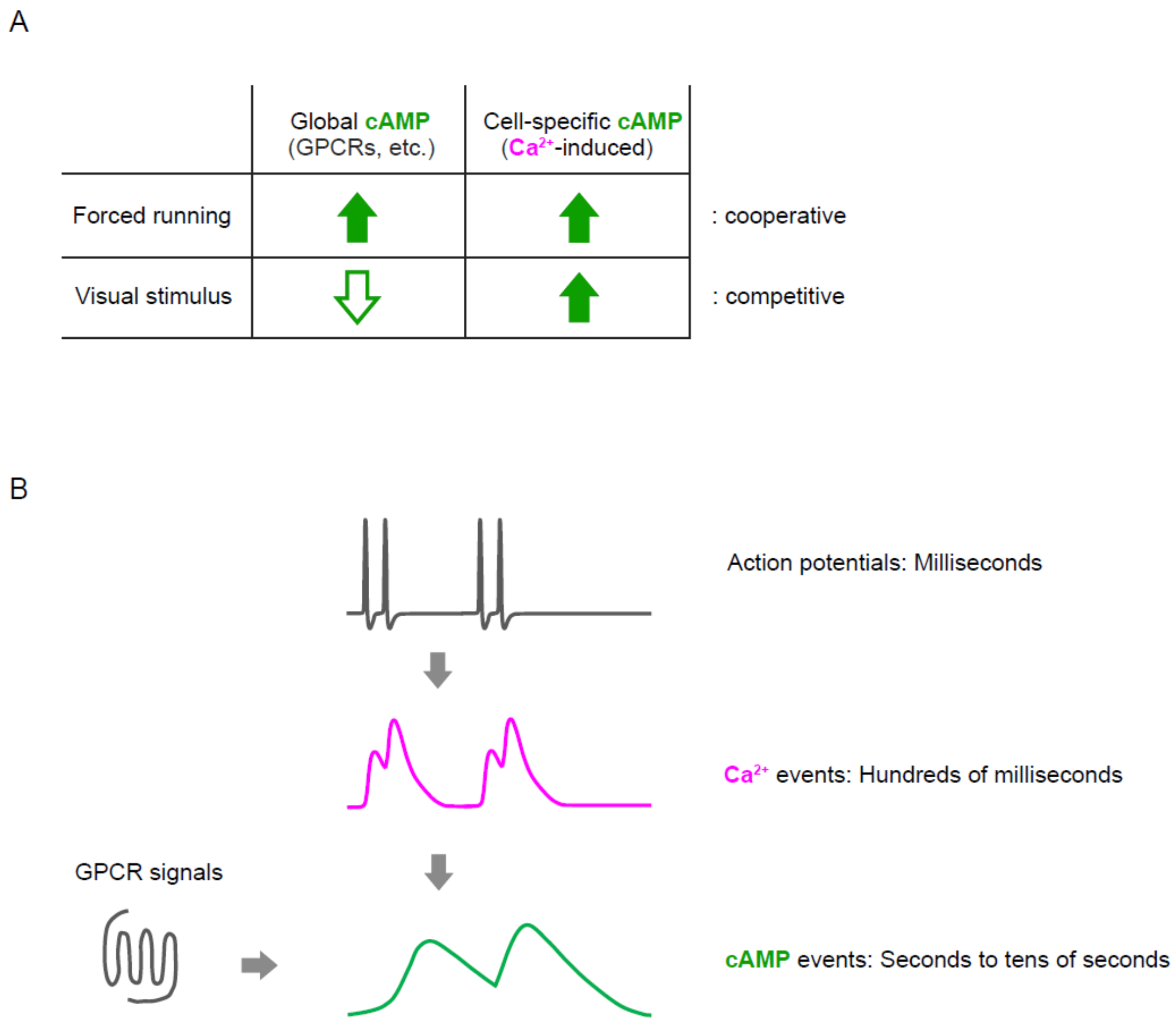


**Figure S9. Summary diagram, related to Discussion**

**(A)** Directions of somatic cAMP change during forced running and visual stimulus under the condition of this study.

**(B)** Multiple upstream of cAMP. The information flows from action potentials to $Ca^{2+}$ events by voltage-dependent calcium channels in individual cells. The information encoded in $Ca^{2+}$ events then flows to cAMP events through $Ca^{2+}$-dependent adenylate cyclases. Because the timescales of action potentials, $Ca^{2+}$ events, and cAMP events are on the order of milliseconds, hundreds of milliseconds, and seconds to tens of seconds, respectively, multiple electrical events are integrated and stored for a longer timescale as cAMP transients through $Ca^{2+}$ signaling. Moreover,

GPCRs can bidirectionally change cAMP levels through G proteins. Overall, information encoded in action potentials, $Ca^{2+}$, and GPCRs signaling is integrated and stored for a longer timescale as cAMP transients.

### Supplementary Movie

**Movie S1. Two-photon mesoscale RCaMP3 imaging in L5, related to Figure 3**

Fast-scanning high optical invariant two-photon microscopy (FASHIO-2PM) for RCaMP3 imaging from L5 neurons in an awake condition. The size of the imaging area is 3.0 x 3.0 $mm^2$ (2,048 x 2048 pixels), acquired at 7.5 Hz.

**Movie S2. *In vivo* dual-color imaging for $Ca^{2+}$ and cAMP during visual stimulation, related to Figure 4**

The mouse was presented with moving grating in eight directions to the contralateral eye in awake condition. Sequential excitation at 940 nm and 1,040 nm was used for dual-color imaging of RCaMP3 (magenta) and cAMPinG1-ST (green). Three optical planes spaced 30 μm apart were imaged at 3.4 Hz per plane using a piezo objective scanner. The size of the imaging area is 339 × 339 $\mu m^2$ (512 × 512 pixels).

**Supplementary Table 1. Comparison of cAMP sensors *in vitro*, related to Figure 1**

| Name | Type | This study | | Literature values | | Ref. |
|---|---|---|---|---|---|---|
| | | ΔF/F | $K_d$ (μM) | ΔF/F | $K_d$ (μM) | |
| Flamindo2 | Single-FP, Green | -0.80 | 2.5 | -0.75 | 3.2 | Odaka et al., PLoS One. 2014 |
| R1α #7 | FRET | - | - | 0.38 | 0.037 | Ohta et al., ACS Chem Biol. 2016 |
| Pink Flamindo | Single-FP, Red | - | - | 4.2 | 7.2 | Harada et al., Sci Rep. 2017 |
| R-FlincA | Single-FP, Red | - | - | 6.0 | 0.30 | Ohta et al., Sci Rep. 2018 |
| cAMPFIRE-H | FRET | - | - | - | 0.38 | Crystian et al., Nat Methods. 2022 |
| gCarvi | Single-FP, Green | 2.2 | 6.5 | 1.5 | 2.0 | Kawata et al., PNAS. 2022 |
| G-Flamp1 | Single-FP, Green | 7.5 | 0.87 | 13.4 | 2.17 | Wang et al., Nat Comm. 2022 |
| G-Flamp2 | Single-FP, Green | - | - | 20.0 | 1.9 | Liu et al., Front Pharmacol. 2022 |
| cAMPinG1 | Single-FP, Green | 10.6 | 0.18 | - | - | This study |

-, not measured or described.
